# Emergence of networks of shared restriction-modification systems in phage-bacteria ecosystems

**DOI:** 10.1101/2021.10.14.464203

**Authors:** Rasmus Skytte Eriksen, Nitish Malhotra, Aswin Sai Narain Seshasayee, Kim Sneppen, Sandeep Krishna

## Abstract

Restriction-modification (RM) systems are the most ubiquitous bacterial defense system against bacteriophages and an important part of controlling phage predation. Using genomic sequence data, we show that RM systems are often shared among bacterial strains in a structured way. Examining the network of interconnections between bacterial strains within each genus, we find that in many genera strains share more RM systems than expected from a random network. We also find that many genera have a larger than expected number of bacterial strains with unique RM systems. We use population dynamics models of closed and open phage-bacteria ecosystems to qualitatively understand the selection pressures that could lead to these non-random network structures with enhanced overlap or uniqueness. In our models we find that the phages impose a pressure that favours bacteria with more RM systems, and more overlap of RM systems with other strains, but in bacteria dominated states this is opposed by the increased cost to the growth rate of these bacteria. Similar to what we observe in the genome data, we find that two distinct bacterial strategies emerge – strains either have a larger overlap than expected, or they have more unique RM systems than one expects from a null model. The former strategy appears to dominate when the repertoire of available RM systems is smaller but the average number of RM systems per strain is larger.

## 2 Introduction

The lives of bacteria can be harsh, with several species competing for limited resources in an environment that is often toxic or which contains other stressors. In addition to these already bleak conditions, bacteria must also defend against a ubiquitous predator - the bacteriophage - which wreaks havoc on bacterial populations. These bacterial viruses outnumber bacteria 10 to 1 and are a major contributor to bacterial death, for instance in the ocean, where phages are responsible for ∼20% of bacterial deaths[1–3]. As a consequence, bacteria have evolved strategies for phage evasion and defence to combat the phage predation, while the phages in turn have evolved counter strategies. This co-evolutionary “arms-race” between phages and bacteria is likely a major selective force shaping the bacterial genome[4–7].

One bacterial defence mechanism is the restriction-modification (RM) system, which is the focus of this paper. RM systems constitute a simple mechanism for a bacterium to distinguish self from non-self. An RM system consists of two parts: a methyltransferase, which methylates (modifies) specific recognition sites in the genome, and an endonuclease, which cleaves (restricts) the genome at the recognition sites when they are not methylated. When a phage infects the bacterium, its genetic material will then be restricted by the RM systems unless it is methylated at the corresponding recognition sites. In rare cases, the phage genome will, by chance, become methylated by the methyltransferase at all of its recognition sites before the endonuclease of the RM system acts. When this happens, the phage genome is said to have “escaped” restriction, and can now freely continue its infection cycle inside the bacterium, eventually leading to lysis and the release of new phage particles, all of which are now methylated at the recognition sites (schematically depicted in Fig. 1). Thus, RM systems are not perfect defences, with the probability of escaping ranging from ∼10^−2^ to 10^−8^ [8–11]. RM systems are remarkably widespread among bacteria, being present in ∼90% of all bacterial strains[12] and occupying on average ∼0.3% of bacterial genomes[7]. However, since the protection is not perfect, bacteria seemingly need to invest in carrying several distinct RM systems[12].

**Figure 1:**
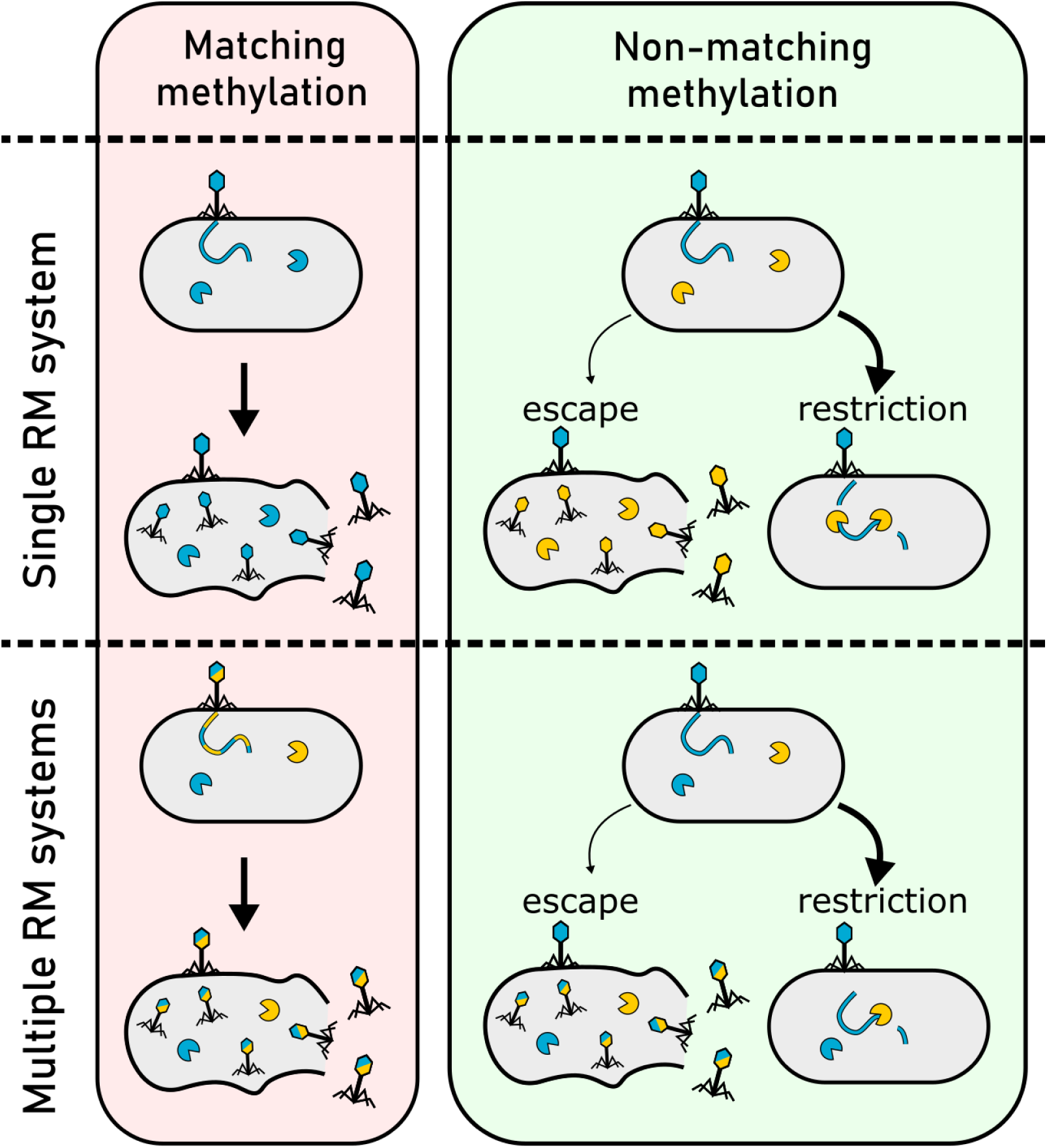
The RM defence mechanism. When a phage infects a bacterium with an RM system, by which it has previously been methylated (indicated by the matching color), then it will be invisible to the RM system and will lyse the host, producing a burst of offspring phage. If the phage lacks the appropriate methylations, the RM system will, a majority of the time, restrict the phage and thereby prevent it from replicating. However, a phage will sometimes escape the restriction and successfully replicate despite lacking the appropriate methylations, and thus produce offspring that carry the methylation pattern that protects it from the host RM system. With multiple RM systems, the phage must carry the combined methylation pattern against *all* RM systems in the host to reliably infect and lyse the host. If it does not, the RM systems which the phage is not protected against will each attempt to restrict the phage. The probability to escape restriction is then dependent on the efficacy of each of these RM systems.

Bacteria have mechanisms for sharing their genetic material between each other (e.g. transduction and conjugation) and as a result, a given RM system is not likely to be unique to its bacterial strain but may be present in other bacterial strains in the surrounding environment. Indeed, sequencing data of bacteria of the same genus reveals that the composition of RM systems within these strains share many common RM systems between each other[13, 14]. In Fig. 1 we show schematically what the sharing or *overlap* of RM systems between related bacterial strains means for their ability to defend against the different epigenetic phage variants. A phage infecting a particular bacterium may already be methylated against some of the RM systems present in that bacterium, rendering these RM systems ineffective. However, by sharing RM systems between related bacterial strains, the bacteria, on average, increase the number of effective RM systems against any given phage which under some conditions may improve their chances of surviving a phage encounter. We do not yet understand under which conditions such sharing or overlap will be advantageous, and under which it will be disadvantageous for bacterial strains.

An ecosystem with multiple bacterial strains with a diverse collection of RM systems can be thought of as a *network* of strains interconnected by the RM systems they share. Our main interest in this paper is to understand how such networks of bacterial strains and their RM systems behave and how they are shaped by evolution: Do these networks have non-random structures? If so, how does the structure dictate the population dynamics? Relatedly, what kind of selection pressures lead to the emergence of these non-random structures? For instance, do certain network structures enhance co-existence of strains or increase the net biomass of the ecosystem?

In this paper, we first examine, in section 3, the RM systems present in publicly available bacterial genomes in the NCBI RefSeq database[15]. Specifically, across 42 bacterial genera, we analyze overall features of RM system use, such as the distribution of the number of RM systems and the degree to which RM systems are shared across strains. In section 4, we then examine the structures of the networks formed by bacterial strains within each genus and the RM systems they contain. We compare these networks to ensembles of randomized networks obtained from a suitable null model, and demonstrate that the real networks often have a higher than expected sharing of RM systems, and many also exhibit a larger than expected number of bacterial strains with unique RM systems. Then, in section 5, we develop a population dynamics model that extends previous approaches to include the sharing of RM systems among bacteria. The model allows us to examine the dynamical effects of RM system sharing in closed and open phage-bacteria ecosystems, and thereby to qualitatively understand what kind of selection pressures may result in the emergence of these non-random network structures. We find in these models that two kinds of strategies seem to work well for bacterial strains - they either have multiple RM systems which they share with other strains, or fewer but unique RM systems - signatures of which are also found in our analysis of bacterial genomes.

## 3 Distribution of RM systems from genome sequencing data

Our data set consists of the complete genomes of 1417 unique bacterial strains across 42 genera which we have analyzed for the presence and absence of 333 different type II RM systems (see Methods and section S1 of the supplement).

Fig. 2 shows the distribution of RM systems across our sequenced genera in two different ways.

**Figure 2:**
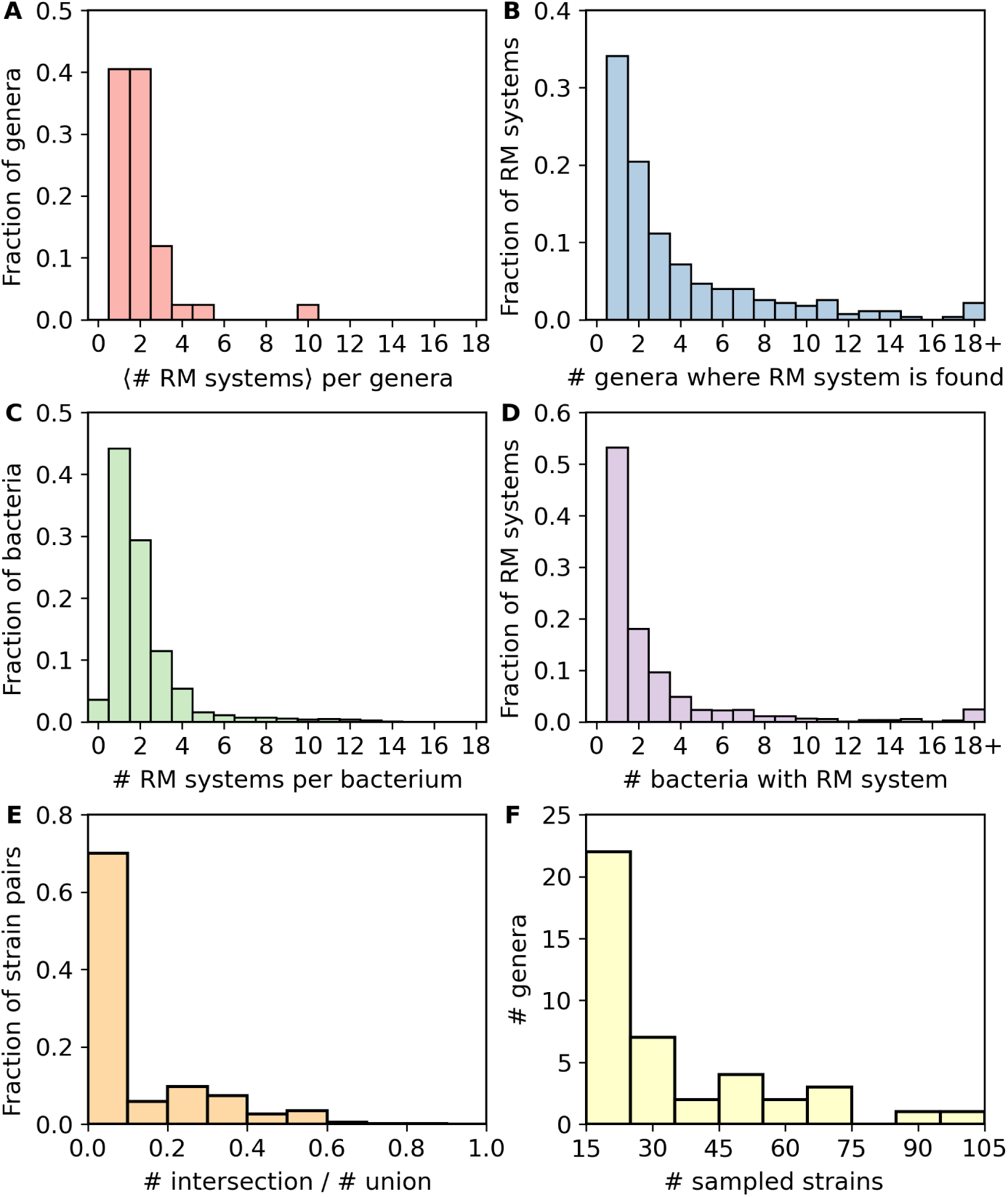
Analysis of RM system distribution in the data set consisting of 279 unique RM systems and 1417 unique bacterial strains (see Methods). From this data, we can compute: (i) for each genus, the presence-absence matrix whose elements, 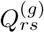 are assigned 1 if RM system *r* is found in bacterial strain *s* of genus *g* and zero otherwise; (ii) a genus-level presence-absence matrix whose elements, *P*_*rg*_, are 1 if RM system *r* is found in genus *g* and zero otherwise (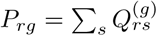, where the sum is over all strains in genus *g*); (iii) the number of strains per genus, *S*(*g*). We first compute for each genus *g* the average number of RM systems in the strains it contains, 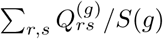 (shown in Supplementary Fig. S2, left panels). The plot in (A) shows the histogram of this data. (B) shows the distribution of genera across RM systems, i.e. it plots Σ_*g*_ *P*_*rg*_. Then, for each genus, *g*, We compute the number of RM systems present in each strain *s* within that genus, i.e. 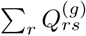. We plot histograms for each genus from this data in Fig. S2 (green bars). (C) shows the average of all these histograms, weighted by *S*(*g*). Then, for each genus, *g*, we compute the number of strains each RM system is found in, i.e., 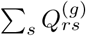 The histograms for each genus from this are shown in Fig. S2 (purple bars). (D) shows the average of all these histograms, weighted by *S*(*g*). Finally, for each genus, *g*, we compute the overlap of RM systems for each strain pair (*i, j*) in that genus. Overlap is defined as the ratio of the number of shared RM system to the total number of distinct RM systems in that pair (see Methods). Histograms of this data for each genus are plotted in Fig. S2 (orange bars). (E) shows an unweighted average of these histograms. (F) shows the distribution of samples across genera, *S*(*g*).

At the genus level, it appears that the majority of genera have few RM systems on average, but with a substantial fraction containing more than 1 RM system on average (Fig. 2A). Fig. 2C shows the same distribution at the level of individual strains within a genus, aggregated across all genera (the contribution of each genus is weighted by the number of samples within our database, see Fig. 2F). As expected, the distribution of RM systems amongst strains has a longer tail than the same distribution at the genus level, due to averaging in the latter. The corresponding distributions for each genus are separately shown in Supplementary Fig. S2. These distributions are unimodal with varying widths. Most are peaked at a small number, 1-2, of RM systems per strain. However there are a few genera such as *Helicobacter* and *Neisseria* which peak at a much higher number of RM systems per strain and contribute to the long tail in figure 2C.

Figs. 2B and 2D show the complementary distributions describing how many genera or strains a given RM system is found within. Supplementary Fig. S2 shows the corresponding distributions within each genus separately. In some genera, such as *Bacillus* or *Staphylococcus*, each RM system is found in a small number of strains. In other genera, the distribution is more skewed, where one or two RM systems are widespread among the bacterial strains in that genus and the remaining RM systems are rare (e.g., see *Mycobacterium* or *Pharobacter*). One also can see genera with a broader distribution, such as *Helicobacter* or *Neisseria*. It is noteworthy that a substantial fraction of RM systems are found both in multiple genera, as well as in multiple strains within a genus. This naturally leads one to ask how often these RM systems are shared across strains. We define RM system “overlap” between a pair of strains to be the ratio of the number of RM systems shared by the two strains to the total number of unique RM systems across both strains, which will be a number ranging from 0 (no overlap) to 1 (complete overlap). Fig. 2E shows the distribution of this measure of overlap, computed first for all pairs within the same genus, and then summed across all genera. Most pairs of bacterial strains do not have any common RM systems. However, there is a substantial fraction of pairs that have a non-zero overlap. One may wonder whether larger overlap occurs only in situations where each member of the pair has a very small number of RM systems with one or two in common (e.g., a pair with 2 RM systems each and 1 in common would give an overlap of 0.33). In fact, in supplementary section S5, we show that there are many pairs with a large number of RM systems each which also have many in common, in particular within *Helicobacter*. Examining the overlap distribution within each genus separately (see Supplementary Fig. S2), we find that most distributions have a high peak at zero overlap, like the aggregated distribution in Fig. 2E, except for *Helicobacter* and *Phaeobacter* where the distribution peaks at an overlap of 0.25.

In summary, the distributions in Fig. 2 and Fig. S2 show that while many bacterial strains have unique RM systems, a substantial fraction share RM systems with other strains of the same genus. This feature was observed in many of the genera we analyzed.

## 4 Non-random networks of interconnected bacterial strains due to shared RM systems

A natural representation of an ecosystem containing multiple strains of bacteria and their RM systems, some of which are shared, is provided by a *network* of interconnected bacterial strains and RM systems. In such a representation, bacterial strains are linked to other strains via RM systems they share. More precisely, a network (or graph) of such a system consists of a set of nodes connected by links. Fig. 3 shows three examples of such a network representation from our database for the genera *Lactococcus, Xanthomonas*, and *Mycoplasma*. These were chosen as examples of bacterial genera with increasing average RM abundance, i.e., networks that range from quite disconnected to well connected. Each bacterial strain is shown as a blue node and each RM system is shown as a red node. If a specific RM system is present within a particular bacterial strain, this is represented by a link connecting those two nodes. Thus, these networks are bipartite; there are no links between two red or two blue nodes, and the “presence-absence matrix” described in Fig. 2 specifies the links that exist between red and blue nodes. In section S3 of Supplementary material we show the network structures for all 42 genera in our database. For genera where each RM system is found in a small number of strains, such as *Bacillus* or *Staphylococcus*, the corresponding networks consist of multiple disconnected pieces. Where one or two RM systems are widespread among the bacterial strains in that genus and the remaining RM systems are rare, such as *Mycobacterium* or *Pharobacter*, the networks have a star-like structure, with many strains connected to one or two RM systems. More densely connected networks are the ones with with a broader distribution, such as *Helicobacter* or *Neisseria*.

**Figure 3:**
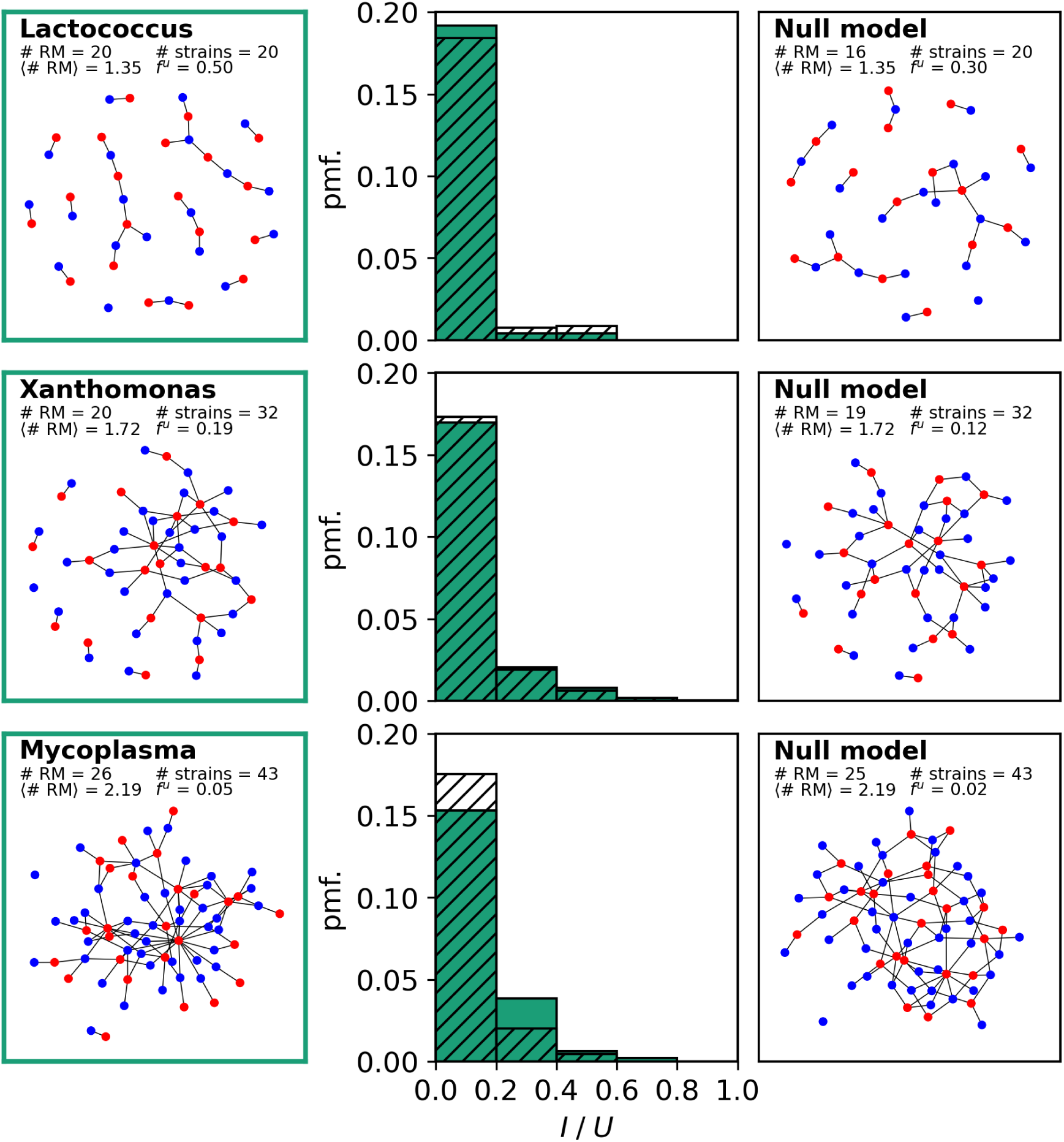
Networks of RM systems. The left column shows bipartite network representations of the distribution of the RM systems among the bacterial strains for three different genera: *Lactococcus, Xanthomonas*, and *Mycoplasma*. Each bacterial strain is shown as a blue node and each RM system is shown as a red node, with links between the two corresponding to the RM system being found within the bacterial strain. The right column shows, for each case, a null model network where the RM systems are chosen randomly (see Methods). The distribution of the overlap in the real (solid green bars) and random network (hatched bars) is shown in the central column.

The network structure will have an effect on the ability of a phage to infect bacterial strains. A genus with a dense network will have strains with many overlapping RM systems, which means that phage successfully infecting one strain will have often have the methylation pattern to avoid some RM systems in a different strain. Whereas disconnected bacterial strains in a sparser network will generally be immune to phage that arise by lysing another strain. We therefore expect such networks to have a non-random structure reflecting the selection pressures that shape the evolution of bacterial strains and their RM systems in the presence of phage. We are most interested in the question: given a repertoire of RM systems and given the constraints that determine the distribution of RM systems across bacterial strains, what are the selection pressures that may lead to enhanced *sharing* of RM systems between strains? Therefore, we have compared each network with a null model, consisting of the same number of bacterial strains and RM systems, with each bacterial strain retaining the same number of RM systems, but where the composition of RM systems is randomized. In other words, for each network, we create an ensemble of randomized networks with the same number of red and blue nodes, where each blue node (bacterial strain) has the same degree (number of links connected to it) as in the real network, but is now connected to randomly chosen red nodes (RM systems).

The right panels in Fig. 3 show one randomized network from the ensemble corresponding to each of the three real networks. There are notable differences in the distribution of RM system overlap between real and random networks, as can be observed visually (notice the relatively fewer red nodes with just one link in the random networks) as well as quantitatively in the histograms in the middle panels. Fig. 4 demonstrates this more rigorously by comparing real networks against an ensemble of 100 random networks, for all the genera. Fig. 4A shows that the average overlap within each genus (as defined previously) differs significantly from the null expectation for many genera. The genera are arranged in order of increasing average RM systems per strain, and interestingly the departure from the null expectation is stronger for genera with a larger average number of RM systems. The visual observation that the real networks seem to have more strains with a unique RM system (red nodes that link to only one blue node) is quantified in Fig. 4B. We define the measure *f* ^*U*^ to be the fraction of bacterial strains that have a unique RM system. We compute *f* ^*U*^ for each real network (i.e., for each genus) and for each network in the corresponding randomized ensemble. Except for a few cases, all genera appear to have significantly more strains with unique RM systems than the random expectation. Note however that this measure is less stable to undersampling than the overlap measure, so we might be overestimating these values (see supplementary figure S1).

**Figure 4:**
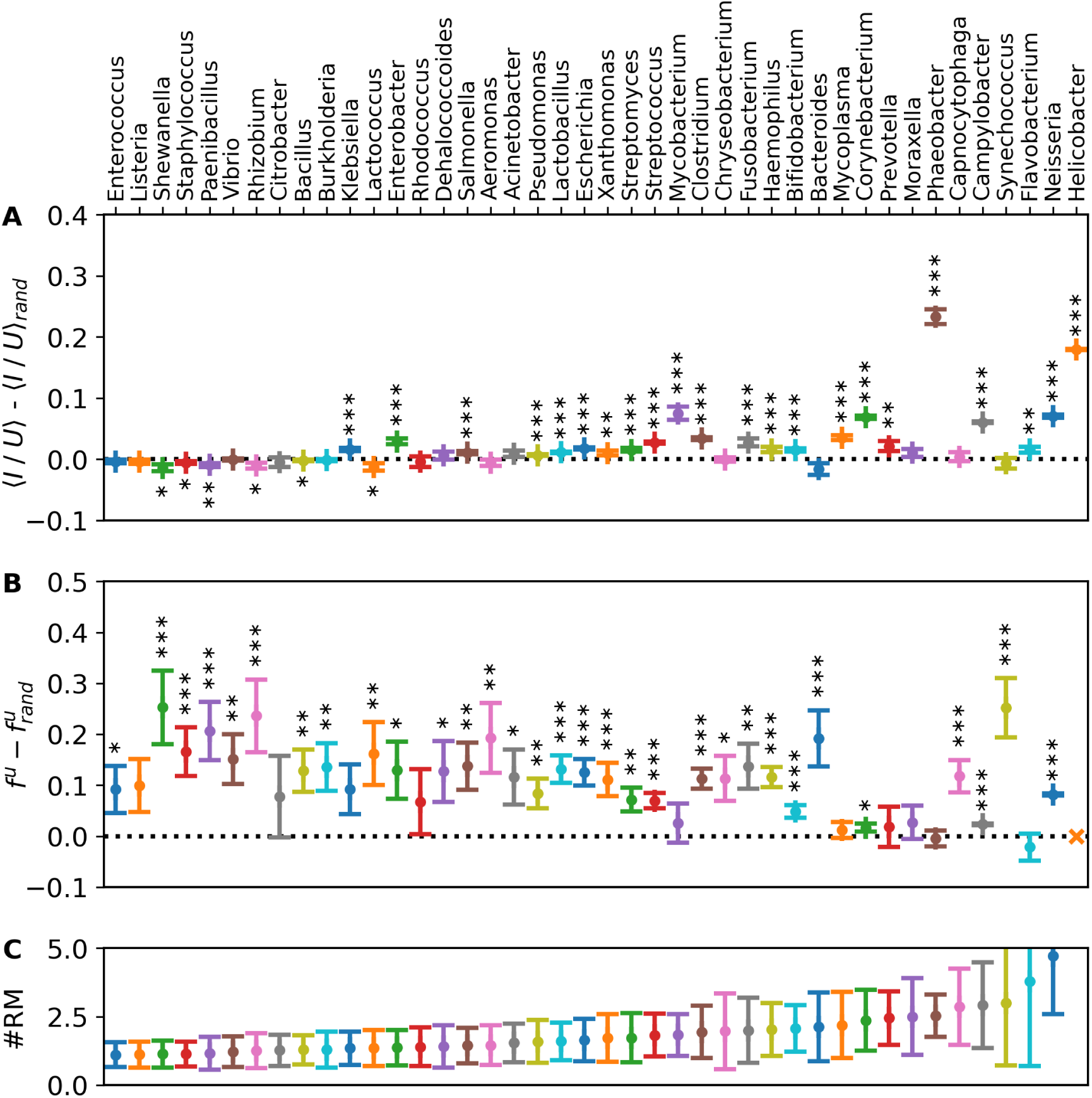
Differences between genome data and the null model. For each of our 42 genera, we have generated 100 samples of the corresponding null model network (see Methods). For each genus, (A) plots the difference in the overlap (averaged over all pairs of strains in that genus) between the real and randomized networks, (B) shows the fraction of bacterial strains with a unique RM system compared to the average of this quantity across the randomized networks for that genus, and (C) plots the average number of RM systems per strain. The y-axis limits were chosen to enhance clarity and do not show the data for *Helicobacter* whose strains have 9.8 RM systems on average, with a standard deviation of 2.9. (A-C) Points indicate averages over the 100 randomized networks, error bars indicate standard deviation, and asterisks denote that the quantities shown are different from zero with a significance level of 95%, 99% and 99.9%, respectively.

We conclude that for a substantial number of genera, selection pressures have led to the emergence of non-random network structures. Specifically, many real strains, even if they have multiple RM systems, more often than expected contain one that is unique to that strain and not shared with other strains. However, there also remains a substantial overlap in RM systems between strains, indicating that any selection pressure to have unique RM systems is not so strong as to eliminate overlap entirely.

In the rest of this paper, we use dynamical models of a simplified phage-bacteria ecosystem with shared RM systems, to obtain a qualitative understanding of the kinds of selection pressures that could lead to such non-random network features.

## 5 Evolution of shared RM systems in mathematical models of phage-bacteria ecosystems

We study the dynamics of phage and bacteria with RM systems in a well-mixed system using equations based on the models of Refs. [16–18], which we have generalized to include sharing of RM systems across bacterial strains (see Methods). Our model implements sharing of RM systems similarly to Ref. [19], which models a spatially extended, not well-mixed, system. Each RM system is assumed to have a corresponding efficacy *ω*_*r*_ and growth rate penalty described by the parameter *γ*_*r*_. For each bacterial strain, its overall growth rate is assumed to be the product of the *γ*_*r*_ of all RM systems it contains, while the overall efficacy of the RM defence can be calculated from the methylation status of the incoming phage (see Methods). First we describe a few simple scenarios to build some intuition about the advantages and disadvantages of sharing RM systems. We term these “closed” ecosystems because the number and kind of strains and RM systems are fixed, although their populations may change with time. We then examine a model of an evolving “open” ecosystem where new strains of bacteria invade, sometimes introducing novel RM systems, and others go extinct. We use these models to understand what selection pressures may lead to networks features similar to those we observed in Sections 3 and 4.

### 5.1 Costs and benefits of overlapping RM systems in simple closed ecosystems

#### 5.1.1 Sharing of RM systems may help increase individual or net biomass

In Fig. 5A we show the case where there is no overlap of RM systems between two bacterial strains. Here, as previously shown[16–18], due to the uniqueness of the RM systems, phage variants have little effect on bacteria that are not from their parent strain. As a consequence, the population density of each bacterial strain is limited primarily by its corresponding phage variant, and all strains reach roughly the same population level irrespective of their intrinsic growth rates. This result generalizes to any number of strains with non-overlapping RM systems, upto the limit imposed by the phage burst size.

**Figure 5:**
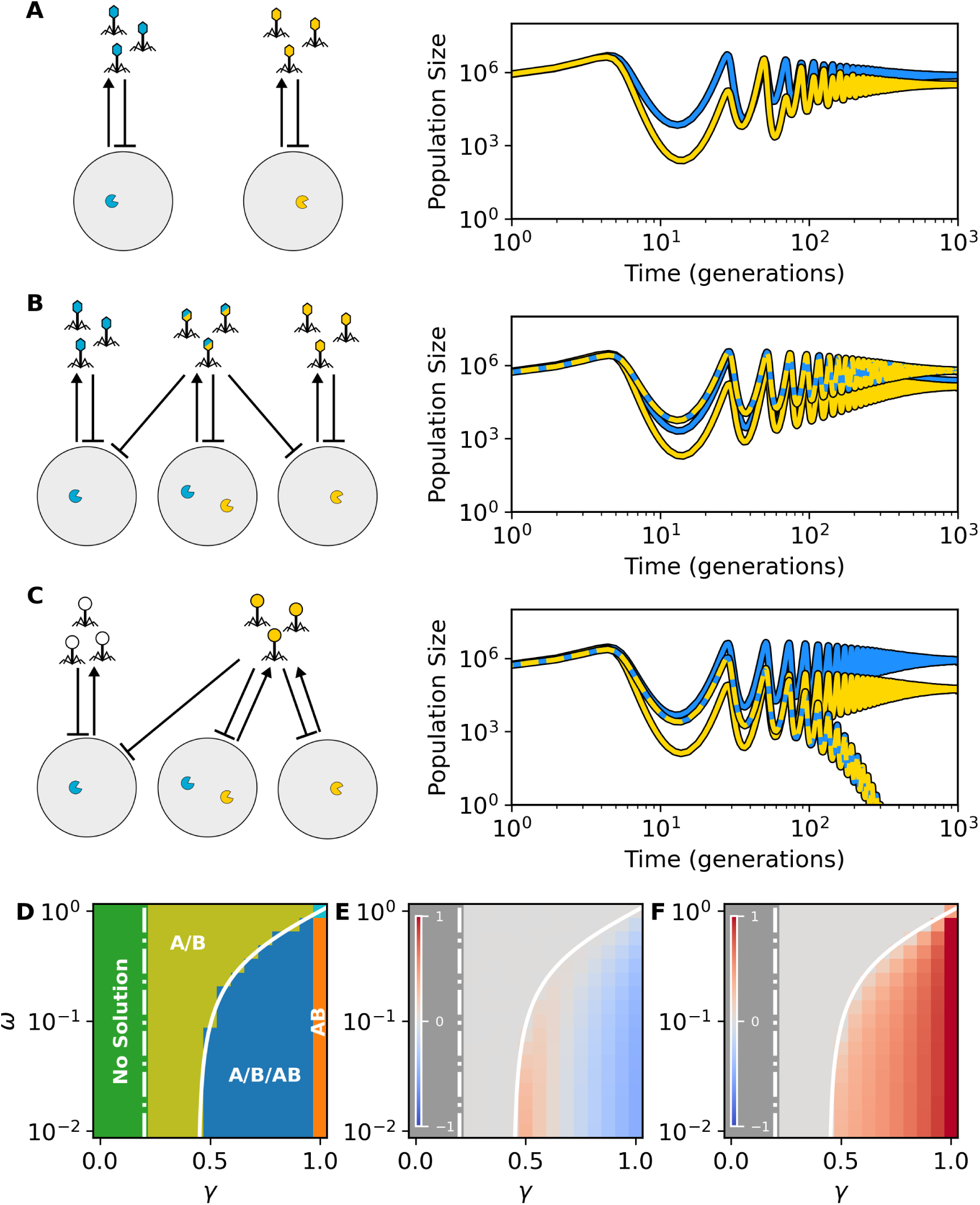
The population dynamics of closed ecosystems. (A) When the bacteria have unique RM systems, all bacterial strains achieve achieve high population densities determined by the efficacy of their respective RM systems. (B) When the RM systems are shared between different hosts, the cross-couplings become more important as some phages will freely attack several hosts. The bacterial strains that are now preyed upon by several phage variants are accordingly less abundant. (C) If a phage has native immunity against an RM system (blue in this case), any host which has invested in the ineffective RM system is now at a disadvantage and likely to be competitively excluded. (D) Regions of possible co-existence solutions for the system in (B) where all RM systems have identical efficacies and costs. (E) Relative gain and loss of biomass for this system compared to the system in (A). (F) Relative fraction of *b*_*AB*_ to *b*_*A*_ + *b*_*B*_ for this system. (A-F) Simulations use default parameters values, except for *β* = 25.

Allowing for overlap between the RM systems changes the situation, even in a simplified example where just two distinct RM systems exist. Here, as shown in Fig. 5B, there are three possible combinations to consider, each with their own epigenetic phage variant. The phage that is methylated at the recognition sites of both RM systems can easily infect all hosts in the system, while the other two phage are primarily limited to their corresponding hosts. Despite the decreased growth rate of the host that carries both RM systems, it may reach a higher level than the other bacteria due to the increased effectiveness of its corresponding phage variant. In other words, due to the overlap, a strain with multiple RM systems may compete better against other strains which have fewer RM systems.

In the special case where all RM systems have identical parameters, i.e. incur the same growthrate penalty (*γ*_*r*_ = *γ*) and have the same efficacy (*ω*_*r*_ = *ω*), we can derive the full solution analytically (see Supplementary section S7.1). In particular, for the system shown in Fig. 5B we can compute when each strain will survive or go extinct, as shown in Fig. 5D. When the growth rate, *γ*, is less than the dilution rate, *α* = 0.2, no bacteria will survive in the long term. When *α < γ < f* (*ω*) (where *f* (*ω*) is a particular function which we derive in Supplementary section S7.1) bacteria with single RM (A or B) begin to survive in the long term but the strain containing both RM systems (AB) is not yet able to overcome its growth rate deficit. When *γ > f* (*ω*) the host with both RM systems now also survives and all three hosts coexist. At *γ* = 1, the bacteria with single RM systems lose their growth advantage and only the bacteria with both RM systems persist. However, when *γ* = *ω* = 1, the system is fully degenerate and all solutions are possible.

Panels E and F in Fig. 5 shows how the biomass changes with the inclusion of the double RM strain (AB). Fig. 5E shows the total biomass in the ecosystem compared to the scenario with the A/B solutions alone. When the growth rate is slightly larger than *f* (*ω*), there is an overall increase in biomass, but as the growth rates increase further, the net biomass decreases due to the presence of the double RM strain. Fig. 5F shows that the biomass of the double RM strain is always greater than the biomass of the single RM strains, which highlights the individual benefit of having several RM systems.

-

In section S7.2 of the supplement, we also consider the high-diversity limit. That is, a scenario where a population of *D* bacterial strains of which *T* are “triplets” that share RM systems in the *A/B/AB* motif (“hierarchical triplets”) while the other *D − T* have unique RM systems. In this case, the range of parameters where the ecosystem can support the double RM system strains is dependent on the number of triplets *T* present in the system. As more and more triplets are added, the range of parameters that support the triplets increase, suggesting that having more RM systems is an invasive strategy. This is further substantiated by considering triplets of the *AB/BC/AC* motif (“looped triplets”), where the range of parameters that support the triplets not only increases faster as *T* increases than for the hierarchical triplets, but also covers a larger range of parameters.

#### 5.1.2 Sharing of RM systems may leave a strain vulnerable to immune phage

However, carrying several RM systems is not always the best strategy, even when they impose only a small cost to the bacterial growth rate. In Fig. 5C we show what happens if the phage develops immunity against one of the RM systems, e.g., by restriction site avoidance[12]. In this example, the phage is immune to the blue RM system, and the strain carrying two RM systems now has an RM system that is ineffective against the phage. This, in turn, prevents the strain from creating a unique epigenetic phage variant. As a consequence, the strain is now in direct competition with the bacterial strain which has not invested in the now ineffective RM system. This competition results in the competitive exclusion of the strain with two RM systems.

Note that for the phage, developing immunity to the RM systems can be a strong selective advantage. In section S6 of the supplement, we show that in the scenario where a non-immune and immune strain of phages compete in the above scenario (i.e. we test the combined scenario of Fig. 5B and 5C), the immune strain will out-compete the non-immune strain if it has no fitness cost associated with the immunity.

### 5.2 Evolution of bacterial networks in an evolving open ecosystem

The above dynamics in model closed ecosystems provides some intuition about the advantages and disadvantages of sharing RM systems across strains, and the kinds of selection pressures that may result in the gain or loss of RM systems. We next consider a more relevant open ecosystem where we periodically add new bacterial strains with potentially new RM systems (and their corresponding phage variants) while removing those that go extinct. As a consequence, we obtain a large open ecosystem which consists of a changing complex interconnected network of bacterial strains, linked by their RM systems. The new strains that are being added have the same average RM abundance as the existing strains but the RM systems are chosen randomly from a set of *K* possible RM systems. In Fig. 6(A-C) we show an example simulation with *K* = 50. Because any particular bacterial strain can have any combination of these RM systems, this corresponds to roughly 10^15^ possible distinct strains. In section S8 of the supplement we compare examples with different values of *K*.

**Figure 6:**
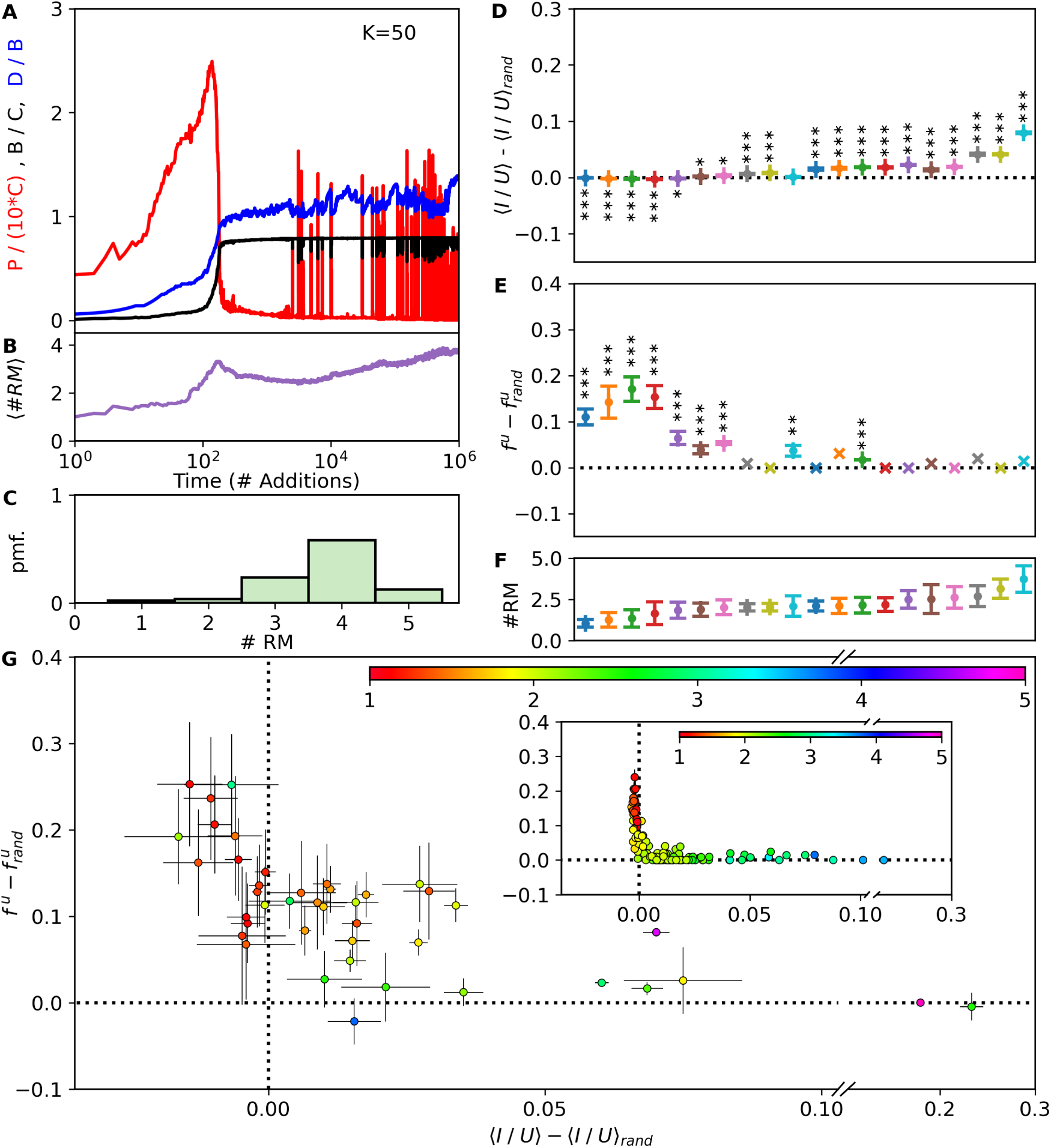
Dynamical link. (A) Example of the population dynamics of open ecosystems. Here, every *T* = 10^3^ generation, we add a new bacterial strain which carries a novel combination of *K* = 50 possible RM systems (see Methods). The plot shows the total bacterial density, *B*, in units of the carrying capacity *C* (black curve), the total phage density *P* (red curve) in units of 10 · *C*, and the diversity *D* (blue curve) in units of the phage burst size *β*. (B) The average number of RM systems per strain #*RM* in the system over time. (C) Distribution of the number RM systems each bacterial strain carries at the end of the simulation. (D-F) We run our simulation with *K* = 800, 400, 200, 10 and 50 and sample the network after the addition of 10^3^, 10^4^, 10^5^, and 10^6^ new strains. For each sampled network, we compute (D) the difference in overlap with 25 realisations of the corresponding randomized network, (E) the difference in fraction of strains with unique RM systems compared to random expectation, and (F) the average number of RM systems per strain for each case. Error bars indicate standard deviation. Asterisks denote significance levels of 95%, 99% and 99.9% respectively. (G) Scatter plot of the two key network characteristics for each genus, obtained by comparing genome data to the null model (from figure 4). The color of each dot indicates the average number of RM systems per strain in that genus. (inset) same as (G) but for the simulations of the open ecosystems (panels D and E). We here include data for 6 repeats of the simulations.

#### 5.2.1 Phage dominated and bacteria dominated states

In the *K* = ∞ limit the system reduces exactly to that studied by Sneppen et al. [17] and Eriksen and Krishna [18] where each strain has a unique RM system and there is zero overlap. Even with a more limited repertoire of available RM systems at finite *K*, and the presence of overlap, some aspects of the dynamics remain the same:

- Early on, competition between bacteria is weak and the phage pressure is high. Due to the relative lack of competition, the diversity (i.e., the number of strains with a non-zero population) increases and the phages pressure decreases. Under these conditions, the ability to defend against phage is more important than the ability to grow fast, therefore bacteria with several RM systems (which typically have a smaller growth-rate) are highly competitive and the average RM abundance increases (see Fig. 6B).
- As the total bacterial population increases and approaches carrying capacity, the ecosystem goes from a phage dominated state to a bacteria dominated state. Bacteria with fewer RM systems (i.e., bacteria with typically higher growth-rates) begin to become competitively viable, which is why the average number of RM systems per strain initially reduces in this phase. However, this does not, in turn, drive the phage pressure back up since the diversity continues to increase and the phage now has a decreasing probability of hitting vulnerable bacteria. In this high diversity, bacteria dominated state there are thus two opposing pressures on the number of RM systems in each strain. For the same reason as seen in the closed ecosystems, there is an advantage in having a larger number of RM systems, and this is opposed by the corresponding cost to the growth rate. The net result is that the average number of RM systems slowly rises and will eventually stabilize when these opposing forces balance (see Supplementary section S8).
- However, the distribution of RM systems per bacterial strain is quite broad (see Fig. 6C). Supplementary section S8 shows that this distribution widens with decreasing diversity of available RM systems, i.e., with decreasing *K*.
- In this bacteria dominated state, the diversity increases slightly beyond the limit set by the phage burst size *β*[17]. Ref. [18] showed that, in the absence of overlap, this happens because of a trade-off between the growth rates of bacterial strains and the strength of their RM systems. In section S8.3 of the supplement, we show that this explanation also applies to this work where we allow for sharing of RM systems across bacterial strains.

#### 5.2.2 Dynamics of overlap of RM systems and intermittent resurgence of the phage

We now focus on the dynamical behaviour that differs from that seen in ref. [18], namely the dynamics of the overlap of RM systems and of the structure of the bacteria-RM networks. In the bacteria dominated phase, the competition between bacteria results in a strong selection for RM systems with small costs (large *γ*_*r*_). This restricts the number of viable RM systems and increases the sharing of RM systems. Thus, the overlap between pairs of strains rises. We find that the number of hierarchical- and looped triplets, discussed in the context of closed ecosystems in section 5.1, increases with time (see Supplement section S8.1). The increase of overlap as time passes in turn increases the chance that the phages are partially protected against the RM systems of the bacterial strains in the ecosystem. When this overlap of RM systems becomes sufficiently large, the phage sometimes experience short resurgences and the dynamics becomes increasingly noisy as can be seen in Fig. 6A. These phage dominated states are not very stable and typically disappear with the next addition of a bacterial strain (see Supplement section S8.2).

#### 5.2.3 Dynamics of the network structure: two distinct bacterial strategies

Fig. 6(D-F) shows an analogue of the plots in Fig 4 for our simulated open ecosystem. Qualitatively similar to what is seen in Fig. 4, we observe that the networks that emerge in our simulations have a larger overlap than random expectation when they contain a larger average number of RM systems. The pattern is more distinct in our simulations - networks that have a lower average number of RM systems more clearly have a lower than expected overlap. We also observe that strains in our evolved networks have more unique RM systems than the random expectation. Here again there is a more distinct pattern in our simulations as a function of the number of RM systems, but the trend is the same. The genome data also shows another interesting pattern when we plot, for each genus, the fraction of strains with unique RM systems compared to random expectation (*f* ^*U*^ − (*f* ^*U*^)_*rand*_) vs the average overlap for that genus compared to random expectation (⟨ *I / U ⟩* - ⟨ *I / U ⟩* _*rand*_). As shown in Fig 6(G), these quantities are anti-correlated, with a higher than expected uniqueness being seen when the overlap is lower than expected and vice versa. This pattern is reproduced in an even starker form in networks from our simulated open ecosystem, as shown in the inset of Fig. 6(G). In the simulated ecosystem, there appear to be two fairly distinct strategies - strains either have a larger overlap than expected but almost the same number of unique RM systems as a random network, or they have the same or lower overlap than the random expectation but have more unique RM systems. In the simulations, the former strategy appears to dominate at later times as the average number of RM systems and the number of hierarchical and looped triplets increases. It also dominates more in our simulations when the repertoire of available RM systems is smaller, i.e., when *K* is smaller. In the genome data there is a similar but weaker correlation with the average number of RM systems, and the two strategies are not so distinctly separated although the trend is qualitatively similar.

## 6 Discussion

In this paper, we investigated the sharing of RM systems between bacterial strains of the same genera, and found large variability across different genera. Depending on the genus of the bacterial strains, the distributions of the RM systems range from cases where the RM systems are shared rather uniformly, to more skewed distributions, where a few RM systems are widespread among the bacterial strains and the remaining RM systems rare. The sharing of RM systems connects the strains and RM systems within each genus into a *network*, whose nodes represent strains and RM systems, and whose links connect RM systems to strains. Mathematically, these networks are bipartite graphs which, we find, have a non-random structure for many genera. Specifically, we observe that genera with a larger average number of RM systems per strain have an RM composition that overlaps more with other strains than expected in a similar class of random networks. We also find that these networks typically have more strains with unique RM systems than the random expectation. The uniqueness and overlap, compared to random expectation, are anti-correlated - genera having a larger average number of RM systems tend to have a higher than expected overlap and a similar uniqueness to random networks, and vice versa.

Extending previous models of ecosystems consisting of a single phage and multiple bacterial strains with RM systems[16–19], allows us to investigate how such patterns of overlap and uniqueness may shape the dynamics of the microbial ecosystem. In our simulations we find that there are opposing selection pressures - the presence of phage favours bacteria with multiple RM systems, but that comes with a cost of lower growth rate, therefore when competition between bacteria is strong an opposing pressure favours strains with less RM systems. In the model we find an even stronger anti-correlation between the overlap of RM systems between strains and the presence of more unique RM systems than the random expectation. Effectively, two distinct strategies appear - bacterial strains either invest in multiple RM systems with an overlap significantly larger than random, or in a larger than expected number of RM systems that are unique and not shared by other strains. In our simulations of an evolving ecosystem, the strains transition from the second strategy to the first at later times when the average number of RM systems is larger. The first strategy, of having larger than expected overlap, also seems to be more prevalent in our model ecosystems when the repertoire of RM systems available to bacteria is smaller. The combination of the phage pressure and the selection pressure towards higher growth rates in the bacteria dominated state drives the transition between strategies.

Existing models of bacteria with RM systems typically lead to only transient dominance by the total phage population over the total bacterial population. When the ecosystem has reached sufficient diversity, the phage can only marginally coexist at very low relative population[17, 18]. Even in a model where RM systems are explicitly allowed to overlap[19], it is only predicted to transiently increase the number of RM systems per bacterium. In the long run, phage population collapses and the number of RM systems will subsequently collapse to about one per coexisting host strain. Notably, in our model, the sharing of RM systems seems to allow the phages to intermittently surpass the density of the bacteria as one observes in real life[1–3].

Our models in many ways represent a limited view of the complex interplay between the bacterial defence systems and phage predators. In particular we primarily focus on cases where a single strain of phage preys on bacteria with an ensemble of RM systems. In the real world, a given host only coexists with about one phage per host in any given environment, but will be exposed to different phages at different times. Our models assume an overall well-mixed approximation to a world that in fact has much fewer phage and bacteria coexisting together at any given time and spatial location. Furthermore, since phages also often exhibit restriction site avoidance, a bacterium will also therefore need to invest in more RM systems than our well-mixed model predicts. Finally, we only consider the influence of a single defence system and thus ignore the effects of CRISPR[20] and abortive infection systems[21]. The inclusion of these defence systems may alleviate some of the need diverse RM systems within each host. Thus, rather than a precise approximation to the interactions occurring in real phage-bacteria ecosystem, our models should be thought of as a way to qualitatively understand the impact of RM system sharing on the population dynamics of phage and bacteria and to provide examples of the kinds of selection pressures which may in turn shape the overlap and uniqueness patterns we observe in the genome data. RM systems are known to serve several functions inside the bacteria (for a review, see Ref. [12]), and these functions have been suggested as an explanation to their abundance in bacteria. Our analysis suggests that interactions with phages alone may impose a net selection pressure that favours increased investment into RM systems and leads to the emergence of networks of shared RM systems between diverse bacterial strains.

## 7 Methods

### 7.1 Distribution of RM systems

Updated sequences and target site information of Type II restriction enzymes and methyltransferases as of March 2020 were obtained from the REBASE database[22]. Sequences of all annotated proteins across complete bacterial genomes as of April 2019 were obtained from NCBI RefSeq database[15]. Sequence homologs for R and M proteins were identified according to Ref. [23]. To simplify the analysis, an RM pair recognising multiple target sequences (e.g. GANT) was considered as a separate pair from another RM pair even if they had overlapping target site sequences (e.g. GACT).

In total, we extracted the presence and absence of 333 RM systems across 12388 bacterial strains. We group these bacterial strains on the level of genera to compare strains which are preyed upon by the same phages. At the level of genera, almost 4 out of 5 bacterial strains have identical composition of RM systems to another strain. For our purposes, we consider two bacteria to be identical if they carry the same set of RM systems.

After filtering, 1021 genera in our data have less than 15 strains with the remaining 42 genera containing containing 1417 strains in total. These strains contain 279 out of the 333 known RM systems. Only these genera with 15 or more samples are included in the analysis.

In summary, our data set includes the presence and absence of 279 RM sequences for 1417 bacterial genomes across 42 different genera, and is represented by a 279 × 1417 “presence-absence matrix” with 1 in the *k*th row and *j*th column if RM system *k* is present in strain *j*, and 0 otherwise. Such presence-absence matrices have been previously studied for *Helicobacter* [13] and *Salmonella*[14] (see Supplementary section S4 for a comparison).

From these presence-absence matrices we can determine: (i) the number of RM systems per strain. (ii) The overlap in RM systems between any two bacterial strains of the same genera. (iii) The network of shared RM systems.

i. The number of RM systems per strain can be readily computed from the presence-absence matrix by summing the RM systems present in each column.
ii. To measure overlap of RM systems, we first describe each bacterial strain *i* by a set *S*_*i*_ containing the IDs of its RM systems. With these list, we can define overlap between strain *i* and strain *j* as the ratio of the number of shared RM systems *I* = |*S*_*i*_ ∩ *S*_*j*_| to the number of unique RM systems *U* = |*S*_*i*_ *∪ S*_*j*_ |across both strains. This measure ranges from 0 for no overlap between the RM systems of the pair, to 1 when the two strains have identical RM systems.
iii. Finally, we construct graphs where RM systems and bacterial strains are represented as nodes and where edges between two nodes signifies the RM system is found in the bacterial strain.

### 7.2 Null model networks

In order to better quantify the network characteristics of the RM distribution, we develop a null model to compare against. For each genus, *g*, we have the presence and absence matrix of the RM systems (i.e. presence-absence matrices). From this data, we then generate corresponding random networks that conform to the following rules:

1. Each bacterial strain in the random network must have the same number of RM systems as the corresponding one in the real network
2. These RM systems, for each strain, are chosen randomly from the *K*_*g*_ available RM systems in the presence-absence matrix (i.e. the strain can contain only RM systems that are present in the sequencing data for that genus).
3. Each randomly generated bacterial strain must be unique in its RM composition (as per our filtering requirement).

Notably, this means that while our null model networks have the same RM abundance distribution (and therefore the same average number of RM systems per bacterial strain), *which* RM systems are present in each strain is randomized. This means that not all of the *K*_*g*_ possible RM systems are necessarily present in the null model networks, and the degree distribution of the RM system nodes is altered.

### 7.3 Population dynamics of a phage-bacteria ecosystem without overlap of RM systems

We base our model on previous models for an ecosystem of *N* bacterial strains which are being preyed upon by a single strain of phage. The bacteria are all valid hosts for the phage, but each bacterial strain carries a unique RM system that protects against the phage. When a phage escapes restriction in a bacterial strain, its offspring will emerge with the methylation corresponding to this “parent” bacterial strain. Therefore, subsequently these phage with this particular methylation pattern can freely infect their parent strain. Eventually, phage will escape restriction from all *N* bacterial strains, giving rise to *N* corresponding epigenetic phage variants. In a well-mixed ecosystem, the equations governing the dynamics of the *N* bacterial strains and the *N* phage variants take the form (see Refs. [16–18]):

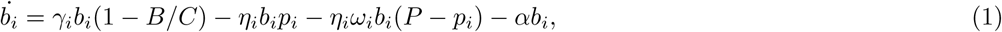

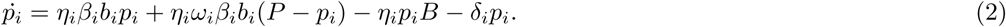

These equations describe the densities of each bacterial strain, *b*_*i*_, and the corresponding epigenetic phage variant, *p*_*i*_. The bacteria grow logistically at a maximal rate *γ*_*i*_ until the total bacteria density, *B* = Σ*b*_*i*_, reaches the carrying capacity *C* = 10^8^ of the ecosystem. In the closed ecosystem models, the value for *C* can be chosen arbitrarily, since the densities can be measured in units of *C* without changing the dynamics. However, in the open ecosystem models, since we remove strains with less than one member, the value is important because 1*/C* controls when strains are considered extinct. The growth rates, *γ*_*i*_, are measured in units of the maximal attainable growth rate of the bacterial strains, and thus are in units of the minimal generation time *τ*. This is the timescale that we measure our parameters relative to. Each phage variant adsorbs to the bacteria at a rate *η*_*i*_ = 10^−8^ in units of 1*/τ*. We run our simulations with a phage burst size *β* = 100 comparable with what is found for real phages[24]. We parametrize the strength of an RM system with the probability *ω*_*i*_ that the phage will bypass the RM system in bacterial strain *i* despite not having the right methylation pattern. Whenever the phage successfully infects a bacterium, *β* new phage particles are produced whose pattern of methylation pattern matches the RM system in the parent bacterium. The model also includes a separate decay rate of the bacteria, *α* = 0.2 *τ* ^−1^, and the phages, *δ*_*i*_ = 0.2 *τ* ^−1^. See table S3 in the supplement for the default parameter values.

### 7.4 Population dynamics with overlapping RM systems

We extend the previous model by considering each bacterial strain to be identified by a unique combination of (one or more) RM systems, some of which may be shared among different bacterial strains. In this case, it is important to keep track of the individual methylation patterns on the epigenetic phage variants and the RM systems of the potential hosts. We achieve this by labelling each RM system with a number *r*, and each bacterial strain by a list of numbers, *S*_*i*_, corresponding to the RM systems it contains. Formally, we define the *i*th bacterial strain by the set of RM systems 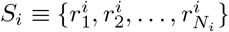 similar to the method in Ref. [19]. Similarly, the *j*th phage variant, which arises by lysing a bacterium of the strain *j*, will have the methylation pattern of its parent bacterium, namely 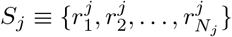.

In addition, where needed we allow phage to have innate immunities against specific RM systems (e.g. via restriction site avoidance[12]). This list of immunities, *I*, is the same for all epigenetic phage variants since it is a property of the the phage itself, independent of its methylation pattern. Combined, this means the efficacy of the RM systems of the *i*th bacterial strain against the *j*th phage variant can be defined by the set of effective RM systems:

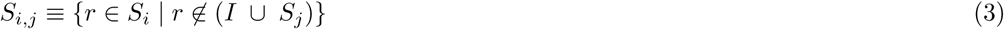

Essentially, this set, *S*_*i,j*_, defines the RM systems in the *i*th bacterial strain that the *j*th phage is *not* immune or epigentically protected against.

Accordingly, the probability of phage *j* escaping the RM systems in bacterial strain *i* becomes:

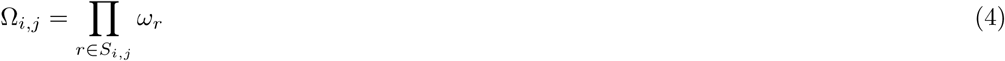

We assume that the cost, to bacterial strains, for having RM systems comes solely from the RM systems in the bacterium. The cost of each RM system, *r*, is encoded in a parameter *γ*_*r*_, such that the growth rate of a bacterial strain is then:

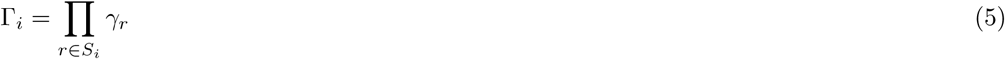

With these definitions, our extended model reads:

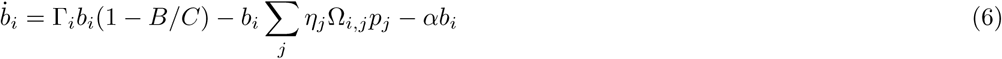

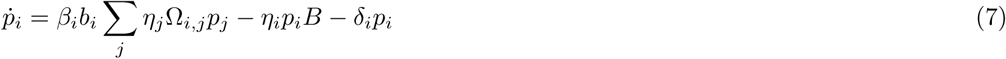

This model is a generalization of the simple model without overlap, from the previous section.

### 7.5 Open ecosystem model

With a fixed number of bacterial strains, the bacterial and phage densities typically reach a steadystate after *O*(10) bacterial generations [17]. However, over even longer timescales, one may expect that new strains of bacteria (with different, unique RM systems) arise by invasion, mutation, or acquisition of RM systems from the environment or other bacteria. As a simplified model of this scenario, we consider an ecosystem consisting of *M* strains of bacteria and the corresponding *M* phage variants, described by the equations above. Periodically, at times *T*, 2*T*, 3*T*, …, we introduce a single bacterium from a new strain with a unique combination of RM systems and a single phage particle with patterns of methylation corresponding to these RM systems. After this addition, we allow the dynamics to proceed for *T* = 10^3^ generations so the populations may reach their steady-state before additions of new strains. Whenever new strains are added, we also remove all bacterial strains and phage variants whose density has fallen below the level corresponding to a single individual.

We initialize the ecosystem with 5 strains each containing a single, randomly chosen, RM system. To allow the number of RM systems per bacterium to change over time, the invading bacteria should resemble the existing bacteria but with sufficient variability to allow for selection of strains. To achieve this, the new bacteria contain *m* random RM systems where *m* is drawn from a Poisson distribution with a mean equal to the median number of RM systems in the current population of bacterial strains.

### 7.6 Choice of parameter values

To completely specify our open ecosystem model, we must describe how the parameters of the initial strains and each new strain are chosen. In Ref. [17], the phage parameters *η, β*, and *δ* were chosen to be the same for all phage variants, whereas *ω*_*i*_ and *γ*_*i*_ were allowed to vary. We do the same (see table S3 for a full list of parameter values), except that we now choose the *ω*_*r*_ and *γ*_*r*_ values for each RM system *r*, instead of each host *i*. The values of *ω*_*r*_ are sampled from a log10-uniform probability distribution between 1 and 10^−4^[8–11]. The *γ*_*r*_ values are chosen independently from a uniform probability distribution between 0.9 and 1[11] corresponding to an average growth rate penalty of 5% per RM system before selection. Our model is constricted by computational complexity in the need to run the simulations for 10^6^ additions of bacteria.

### 7.7 Implementation and data availability

Our analyses and models are implemented partly in MATLAB and partly in python. The code used in our analysis is based on the code[25] used in Ref. [18]. We have made the full code and data available in the online repository located here[26]: https://github.com/RasmusSkytte/OverlappingRestrictionModificationSystems/tree/v1.0

## 8 Funding

R.S.E and K.S. have received funding for this project from the European Research Council (ERC) under the European Union’s Horizon 2020 research and innovation programme under grant agreement No 740704. N.M., A.S.N.S. and S.K. acknowledge support of the Department of Atomic Energy, Government of India, under Project Identification No. RTI 4006. N.M and A.S.N.S are funded by DBT/Wellcome Trust India Alliance Intermediate Fellowship (IA/I/16/2/502711 to A.S.N.S.). S.K. also acknowledges the Simons Foundation (Grant No. 287975) for funding.

## Notes

https://github.com/RasmusSkytte/OverlappingRestrictionModificationSystems

### Competing Interest Statement

The authors have declared no competing interest.

