## Supplement for "Emergence of networks of shared restriction-modification systems in phage-bacteria ecosystems"

### Supplementary Material for: Emergence of networks of shared restriction-modification systems in phage-bacteria ecosystems

Rasmus Skytte Eriksen[1,2], Nitish Malhotra[3], Aswin Sai Narain Seshasayee[3], Kim Sneppen[1] & Sandeep Krishna[4],

[1] Niels Bohr Institute, University of Copenhagen, Copenhagen, Denmark

[2] Statens Serum Institut, Copenhagen, Denmark

[3] National Centre for Biological Sciences, Tata Institute of Fundamental Research, Bangalore, India.

[4] Simons Centre for the Study of Living Machines, National Centre for Biological Sciences, Tata Institute of Fundamental Research, Bangalore, India.

September 30, 2021

#### Contents

**I Data** 2

**S1 List of RM systems** 2

**S2 Network stability and sampling frequency** 6

**S3 Networks of RM systems** 7

**S4 Comparison with previous experiments** 14

**S5 Breakdown of the overlap distribution** 15

**II Model** 16

**S6 Fitness advantage of restriction site avoidance** 16

**S7 Possible solutions to the model equations** 17
- S7.1 Low diversity limit 17
- S7.2 High diversity limit 19
  - S7.2.1 Solution to embedded hierarchical triplets 19
  - S7.2.2 Solution to embedded looped triplets 20
  - S7.2.3 Embedded triplets 21

**S8 Examples of the open-ecosystem** 22
- S8.1 Triplet composition 23
- S8.2 Phage resurgences 23
- S8.3 Extinction curve 24

**S9 Parameter values used in simulations** 25

### Part I
### Data

#### S1 List of RM systems

We extracted 333 RM sequences from the REBASE database[1]. After filtering, the 42 bacterial genera that we include in our analysis contain 279 of these 333 RM sequences. In table S1 we provide the target sequences of the found RM motifs. In table S2 we provide the target sequences not found in the filtered data. Note that we provide the target sequences as: (R target)-(M target).

Table S1: List of the target sequences for the 279 RM systems included in this analysis. The target sequences are reported as (R target) - (M target).

| | | | |
|---|---|---|---|
| 1 | AACNNNNNGTT-AACNNNNNGTT | 41 | CACRAG-CACRAG |
| 2 | AAGCTT-AAGCTT | 42 | CAGCTG-CAGCTG |
| 3 | AAGGAG-AAGGAG | 43 | CAGNNNCTG-CAGNNNCTG |
| 4 | ACACAG-ACACAG | 44 | CAGRAG-CAGRAG |
| 5 | ACCCAC-ACCCAC | 45 | CATATG-CATATG |
| 6 | ACCYAC-ACCYAC | 46 | CATCAC-CATCAC |
| 7 | ACGABGG-ACGABGG | 47 | CATCAG-CATCAG |
| 8 | ACGCAG-ACGCAG | 48 | CATCGAC-CATCGAC |
| 9 | ACGGA-ACGGA | 49 | CATCNAC-CATCNAC |
| 10 | ACGGC-ACGGC | 50 | CATG-CATG |
| 11 | ACGRAG-ACGRAG | 51 | CAYNNNNRTG-CAYNNNNRTG |
| 12 | ACGT-ACGT | 52 | CCAGA-CCAGA |
| 13 | ACNGT-ACNGT | 53 | CCANNNNNNGT-CCANNNNNNGT |
| 14 | ACNNNNNCTCC-ACNNNNNCTCC | 54 | CCANNNNNNTGG-CCANNNNNNTGG |
| 15 | ACRGAG-ACRGAG | 55 | CCANNNNNTGG-CCANNNNNTGG |
| 16 | ACTGG-ACTGG | 56 | CCATC-CCATC |
| 17 | ACTGGG-ACTGGG | 57 | CCATGG-CCATGG |
| 18 | AGATCT-AGATCT | 58 | CCCAAG-CCCAAG |
| 19 | AGCABCC-AGCABCC | 59 | CCCGAG-CCCGAG |
| 20 | AGCACC-AGCACC | 60 | CCCGC-CCCGC |
| 21 | AGCANCC-AGCANCC | 61 | CCCGGG-CCCGGG |
| 22 | AGCCAG-AGCCAG | 62 | CCGC-CCGC |
| 23 | AGCCCA-AGCCCA | 63 | CCGCAG-CCGCAG |
| 24 | AGCT-AGCT | 64 | CCGCGG-CCGCGG |
| 25 | AGGAAT-AGGAAT | 65 | CCGG-CCGG |
| 26 | AGGAG-AGGAG | 66 | CCKAAG-CCKAAG |
| 27 | AGGCCT-AGGCCT | 67 | CCNGG-CCNGG |
| 28 | AGGRAG-AGGRAG | 68 | CCNNGG-CCNNGG |
| 29 | AGTACT-AGTACT | 69 | CCRCTC-CCRCTC |
| 30 | ARCCCC-ARCCCC | 70 | CCRGAG-CCRGAG |
| 31 | ATCGAT-ATCGAT | 71 | CCSGG-CCSGG |
| 32 | ATGAAG-ATGAAG | 72 | CCTC-CCTC |
| 33 | ATGCAT-ATGCAT | 73 | CCTNAGG-CCTNAGG |
| 34 | CAAGNAC-CAAGNAC | 74 | CCTTC-CCTTC |
| 35 | CAARCA-CAARCA | 75 | CCTTGA-CCTTGA |
| 36 | CAATNAG-CAATNAG | 76 | CCTYNA-CCTYNA |
| 37 | CAATTG-CAATTG | 77 | CCWGG-CCWGG |
| 38 | CACGCAG-CACGCAG | 78 | CCWWGG-CCWWGG |
| 39 | CACNCAC-CACNCAC | 79 | CCYCAG-CCYCAG |
| 40 | CACNNNGTG-CACNNNGTG | 80 | CCYGA-CCYGA |

| | | | |
|---|---|---|---|
| 81 | CGAGG-CGAGG | 136 | GACNNNNRTGA-GACNNNNRTGA |
| 82 | CGANNNNNNTGC-CGANNNNNNTGC | 137 | GAGCTC-GAGCTC |
| 83 | CGANNNNNRTAY-CGANNNNNRTAY | 138 | GAGG-GAGG |
| 84 | CGATCG-CGATCG | 139 | GAGGAC-GAGGAC |
| 85 | CGCAGCG-CGCAGCG | 140 | GAGGAG-GAGGAG |
| 86 | CGCG-CGCG | 141 | GAGGC-GAGGC |
| 87 | CGGAAG-CGGAAG | 142 | GAGNNNNNGT-GAGNNNNNGT |
| 88 | CGGCCG-CGGCCG | 143 | GAGNNNNNRTG-GAGNNNNNRTG |
| 89 | CGRGA-CGRGA | 144 | GAGTC-GASTC |
| 90 | CGTACG-CGTACG | 145 | GANGGAG-GANGGAG |
| 91 | CGTARC-CGTARC | 146 | GANTC-GANTC |
| 92 | CGWCG-CGWCG | 147 | GARGAAG-GARGAAG |
| 93 | CNYACAC-CNYACAC | 148 | GASTC-GAGTC |
| 94 | CRAGCAC-CRAGCAC | 149 | GASTC-GASTC |
| 95 | CRTTAA-CRTTAA | 150 | GATAAT-GATAAT |
| 96 | CRTTGAC-CRTTGAC | 151 | GATATC-GATATC |
| 97 | CTAG-CTAG | 152 | GATC-GATC |
| 98 | CTAMRAG-CTAMRAG | 153 | GATCAG-GATCAG |
| 99 | CTATCAV-CTATCAV | 154 | GATCGAG-GATCGAG |
| 100 | CTBVAG-CTBVAG | 155 | GATGC-GATGC |
| 101 | CTCAAT-CTCAAT | 156 | GAYNNNNNRTC-GAYNNNNNRTC |
| 102 | CTCAG-CTCAG | 157 | GCAAAT-GCAAAT |
| 103 | CTCAG-CTSAG | 158 | GCAAGG-GCAAGG |
| 104 | CTCGAG-CTCGAG | 159 | GCAGC-GCAGC |
| 105 | CTCTTC-CTCTTC | 160 | GCAGCC-GCAGCC |
| 106 | CTGAAG-CTGAAG | 161 | GCAGGC-GCAGGC |
| 107 | CTGCAG-CTGCAG | 162 | GCAGT-GCAGT |
| 108 | CTGGAG-CTGGAG | 163 | GCANNNNNNTCC-GCANNNNNNTCC |
| 109 | CTKMAG-CTGAAG | 164 | GCATC-GCATC |
| 110 | CTKMAG-CTKMAG | 165 | GCATGC-GCATGC |
| 111 | CTKMAG-CTRYAG | 166 | GCCGAG-GCCGAG |
| 112 | CTNAG-CTNAG | 167 | GCCGC-GCCGC |
| 113 | CTRYAG-CTRYAG | 168 | GCCGC-GCSGC |
| 114 | CTTAAG-CTTAAG | 169 | GCCGGC-GCCGGC |
| 115 | CTYRAG-CTCGAG | 170 | GCCGNAC-GCCGNAC |
| 116 | CWTCCAG-CWTCCAG | 171 | GCCNNNNNGGC-GCCNNNNNGGC |
| 117 | CYAAANG-CYAAANG | 172 | GCCTA-GCCTA |
| 118 | CYANNNNNNNTTC-CYANNNNNNNTTC | 173 | GCGATG-GCGATG |
| 119 | CYCGRG-CYCGRG | 174 | GCGC-GCGC |
| 120 | GAAABCC-GAAABCC | 175 | GCGCGC-GCGCGC |
| 121 | GAACNNNNNNTCC-GAACNNNNNNTCC | 176 | GCGGAG-GCGGAG |
| 122 | GAAGA-GAAGA | 177 | GCGGCCGC-GCGGCCGC |
| 123 | GAAGAC-GAAGAC | 178 | GCGGRAG-GCGGRAG |
| 124 | GAAGGC-GAAGGC | 179 | GCGTA-GCGTA |
| 125 | GAANCAG-GAANCAG | 180 | GCMGAAG-GCMGAAG |
| 126 | GAATTC-GAATTC | 181 | GCNGC-GCNGC |
| 127 | GACATC-GACATC | 182 | GCNNGC-GCNNGC |
| 128 | GACCAC-GACCAC | 183 | GCRGAAG-GCRGAAG |
| 129 | GACGAG-GACGAG | 184 | GCSGC-GCCGC |
| 130 | GACGC-GACGC | 185 | GCSGC-GCNGC |
| 131 | GACGCA-GACGCA | 186 | GCSGC-GCSGC |
| 132 | GACGTC-GACGTC | 187 | GCTAAT-GCTAAT |
| 133 | GACNNNGTC-GACNNNGTC | 188 | GCTAGC-GCTAGC |
| 134 | GACNNNNGTC-GACNNNNGTC | 189 | GCTNAGC-GCTNAGC |
| 135 | GACNNNNNTGA-GACNNNNNTGA | 190 | GCVGAG-GCVGAG |

Table S1 (cont.)

| | | | |
|---|---|---|---|
| 191 | GCWGC-GCAGC | 236 | GTCTC-GTCTC |
| 192 | GCWGC-GCNGC | 237 | GTGCAG-GTGCAG |
| 193 | GCWGC-GCWGC | 238 | GTGGNAG-GTGGNAG |
| 194 | GDGCHC-GDGCHC | 239 | GTMKAC-GTMKAC |
| 195 | GGACY-GGACY | 240 | GTNNAC-GTNNAC |
| 196 | GGAGGC-GGAGGC | 241 | GTRAAG-GTRAAG |
| 197 | GGANNAG-GGANNAG | 242 | GTSAC-GTSAC |
| 198 | GGARGA-GGARGA | 243 | GTTAAC-GTTAAC |
| 199 | GGATC-GGATC | 244 | GTTAAT-GTTAAT |
| 200 | GGATCC-GGATCC | 245 | GTTCNAC-GTTCNAC |
| 201 | GGATG-GGATG | 246 | GTYGGAG-GTYGGAG |
| 202 | GGATTY-GGATTY | 247 | GTYRAC-GTYRAC |
| 203 | GGCC-GGCC | 248 | GWGCWC-GWGCWC |
| 204 | GGCGAG-GGCGAG | 249 | MTCGAK-CTCGAG |
| 205 | GGCGCA-GGCGCA | 250 | MTCGAK-MTCGAK |
| 206 | GGCGCC-GGCGCC | 251 | RAATTY-RAATTY |
| 207 | GGGAC-GGGAC | 252 | RCCGGY-RCCGGB |
| 208 | GGGTDA-GGGTDA | 253 | RCCGGY-RCCGGY |
| 209 | GGNCC-GGNCC | 254 | RGAAAGR-RGAAAGR |
| 210 | GGNNCC-GGNNCC | 255 | RGATCY-RGATCY |
| 211 | GGRCA-GGRCA | 256 | RGCGCY-RGCGCY |
| 212 | GGRCAG-GGRCAG | 257 | RGGNCCY-RGGNCCY |
| 213 | GGTACC-GGTACC | 258 | RGGWCCY-RGGWCCY |
| 214 | GGTCTC-GGTCTC | 259 | RTAAAYG-RTAAAYG |
| 215 | GGTGA-GGTGA | 260 | RTCAGG-RTCAGG |
| 216 | GGTNACC-GGTNACC | 261 | SAGCTS-SAGCTS |
| 217 | GGWCC-GGWCC | 262 | TAGGAG-TAGGAG |
| 218 | GGWCNA-GGWCNA | 263 | TANAAG-TANAAG |
| 219 | GGWTAA-GGWTAA | 264 | TCANNNNNNTRG-TCANNNNNNTRG |
| 220 | GGYGAB-GGYGAB | 265 | TCCGGA-TCCGGA |
| 221 | GGYRCC-GGYRCC | 266 | TCGA-TCGA |
| 222 | GMCCC-GCCCC | 267 | TCGAG-TCGAG |
| 223 | GNGAAAY-GNGAAAY | 268 | TCGCGA-TCGCGA |
| 224 | GRAGCAG-GRAGCAG | 269 | TCNGA-TCNGA |
| 225 | GRCGYC-GRCGYC | 270 | TCNNGA-TCNNGA |
| 226 | GRGCRAC-GRGCRAC | 271 | TCTAGA-TCTAGA |
| 227 | GRGCYC-GRGCYC | 272 | TCTGG-TCTGG |
| 228 | GRGGAAG-GRGGAAG | 273 | TGATCA-TGATCA |
| 229 | GTAAG-GTAAG | 274 | TGCA-TGCA |
| 230 | GTAATC-GTAATC | 275 | TGGCCA-TGGCCA |
| 231 | GTAC-GTAC | 276 | TGRYCA-TGRYCA |
| 232 | GTATCC-GTATCC | 277 | TTCGAA-TTCGAA |
| 233 | GTATNAC-GTATNAC | 278 | WCCGGW-WCCGGW |
| 234 | GTCGAC-GTCGAC | 279 | YGGCCR-YGGCCR |
| 235 | GTCGAC-RTCGAY | | |

Table S1 (cont.)

Table S2: List of the target sequences for the 54 RM systems not found in the 42 filtered genera. The target sequences are reported as (R target) - (M target).

| | | | |
|---|---|---|---|
| 1 | AAAAGRG-AAAAGRG | 28 | GAGAYGT-GAGAYGT |
| 2 | AACGTT-AACGTT | 29 | GASTC-GANTC |
| 3 | ACGCGT-ACGCGT | 30 | GCAGC-GCWGC |
| 4 | ACRCAG-ACRCAG | 31 | GCTCCA-GCTCCA |
| 5 | ACRYGT-ACRYGT | 32 | GGACGAC-GGACGAC |
| 6 | ATGCAC-ATGCAC | 33 | GGCCGGCC-GGCCGGCC |
| 7 | ATTAAT-ATTAAT | 34 | GGCGGAG-GGCGGAG |
| 8 | ATTTAAAT-ATTTAAAT | 35 | GGGCCC-GGGCCC |
| 9 | CACGTG-CACGTG | 36 | GGNCC-GGWCC |
| 10 | CAGAAG-CAGAAG | 37 | GGNNCC-GRCGYC |
| 11 | CAGCAG-CAGCAG | 38 | GGNNCC-RGCGCY |
| 12 | CCCGT-CCCGT | 39 | GGWCC-GGNCC |
| 13 | CCGGNAG-CCGGNAG | 40 | GNAAYG-GNAAYG |
| 14 | CCRYGG-CCRYGG | 41 | GNGCAAC-GNGCAAC |
| 15 | CCTAGG-CCTAGG | 42 | GRACGAC-GRACGAC |
| 16 | CCTNNNNNAGG-CCTNNNNNAGG | 43 | GRCGYC-GGNNCC |
| 17 | CCWGG-CCNGG | 44 | GTATAC-GTATAC |
| 18 | CGACCAG-CGACCAG | 45 | GTGCAC-GTGCAC |
| 19 | CGRKA-CGRKA | 46 | RCATGY-RCATGY |
| 20 | CMGCKG-CMGCKG | 47 | RGCGCY-GGNNCC |
| 21 | CRCCGGYG-CRCCGGYG | 48 | SAGCTS-CAGCTG |
| 22 | CRRTAAG-CRRTAAG | 49 | SAGCTS-GAGCTC |
| 23 | CTGGAG-CTBVAG | 50 | TACCNAG-TACCNAG |
| 24 | CTKMAG-CTGCAG | 51 | TGCGCA-TGCGCA |
| 25 | CTSAG-CTSAG | 52 | TTAA-TTAA |
| 26 | CTYRAG-CTYRAG | 53 | TTTAAA-TTTAAA |
| 27 | GAAGNAG-GAAGNAG | 54 | YACGTR-YACGTR |

#### S2 Network stability and sampling frequency

The number of sampled strains varies considerably across the different genera. As we noted in the main text, only 42 out of 1063 genera have more than 15 samples. For these 42 genera, the number of samples also vary substantially (see Fig. 2F in the main text). To test whether the large heterogeneity we observe in the network structures (see figure S2) is determined by the differences in sampling between the strains, we run a bootstrapping test on the RM system networks of *Lactococcus*, *Mycoplasma*, and *Xanthomonas*, which are used as examples in Fig. 3.

For the bootstrapping test, we sub-sample our data at frequencies $10\%, 20\%, 30\%, \dots 100\%$. For example, this means that for a sampling frequency of 20%, we randomly choose 20 percent of the samples in our data of a given genus and construct the network based on this subset of samples. We repeat this sampling 100 times for each sampling frequency and genus.

This allows us to measure how the average RM abundance and the overlap metric changes with the sampling frequency of the genus. In Fig. S1 we show that the network overlap does not change significantly for the three sub-sampled genera, whereas the fraction of strains with unique RM systems is more susceptible to the number of samples.

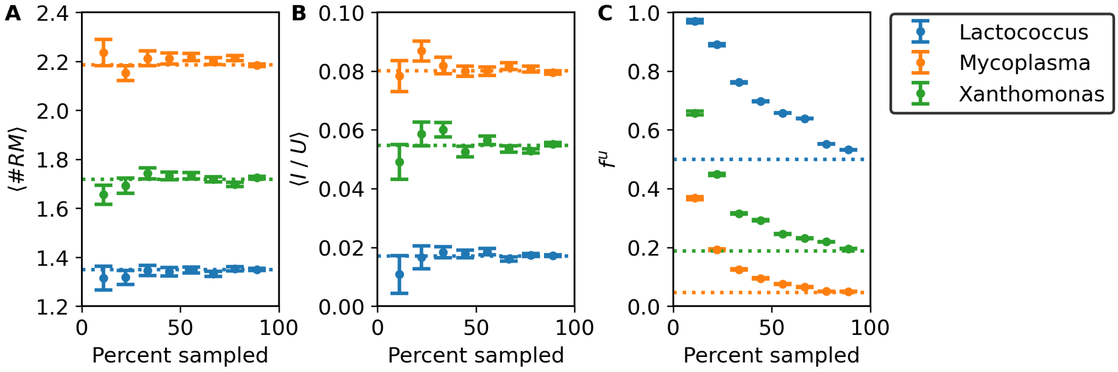


Figure S1: Sampling frequency and metrics. We show as a function of the sampling frequency (A) the average number of RM systems per bacterium. (B) the average overlap: $I\ /\ U$, and (C) the fraction of strains with unique RM systems: $f^u$. (A-C) the information is obtained by bootstrapping our data for 3 different genera: *Lactococcus*, *Mycoplasma*, and *Xanthomonas*. We run the bootstrapping 100 times for sampling frequencies of 90% or below and include the fully sampled genus as a reference (dashed lines). The error bars correspond to the standard error.

### S3 Networks of RM systems

In total, we consider the distribution of RM systems among 43 different genera. In the main text we show examples of the networks of shared RM systems for 3 of these genera, and here in this section we include networks for all 43 analyzed genera (see figure S2).

Previous publications for *Halobacterium* Roer et al. [2] and *Salmonella* Fullmer et al. [3] have made the data available in a form which we could use to plot the distributions they obtained which can be compared to the ones produced by our data in Fig. S2. We do this in the next section S4.

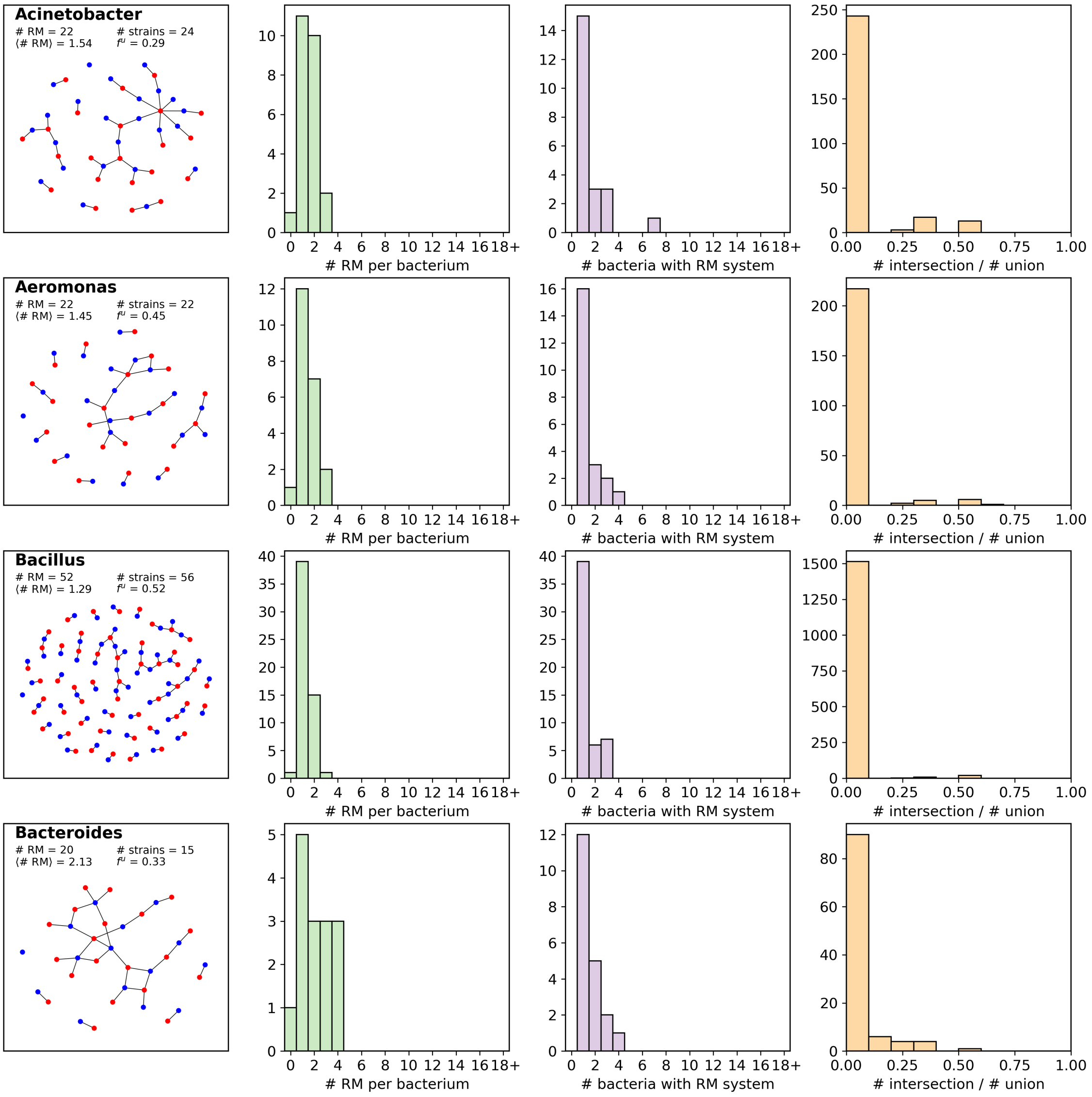


Figure S2: All networks and distributions of RM systems. Our full data set contains 42 different genera with at least 15 sampled strains (after filtering). Here we show the network of RM systems for each of these genera (see Methods). We further show the histograms corresponding to Fig. 2C-E for each genus. Figure continues below.

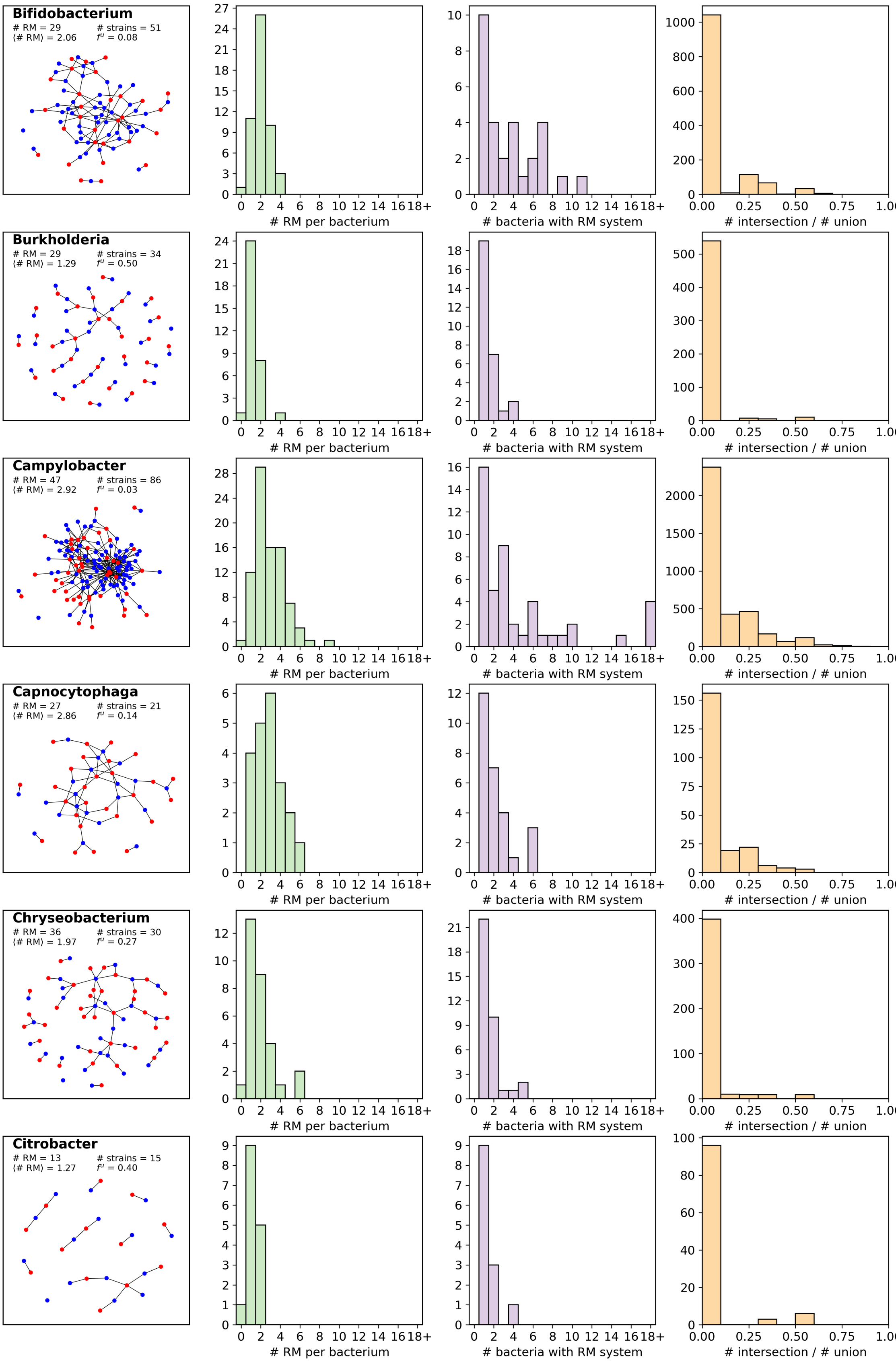


Figure S2 (cont.)

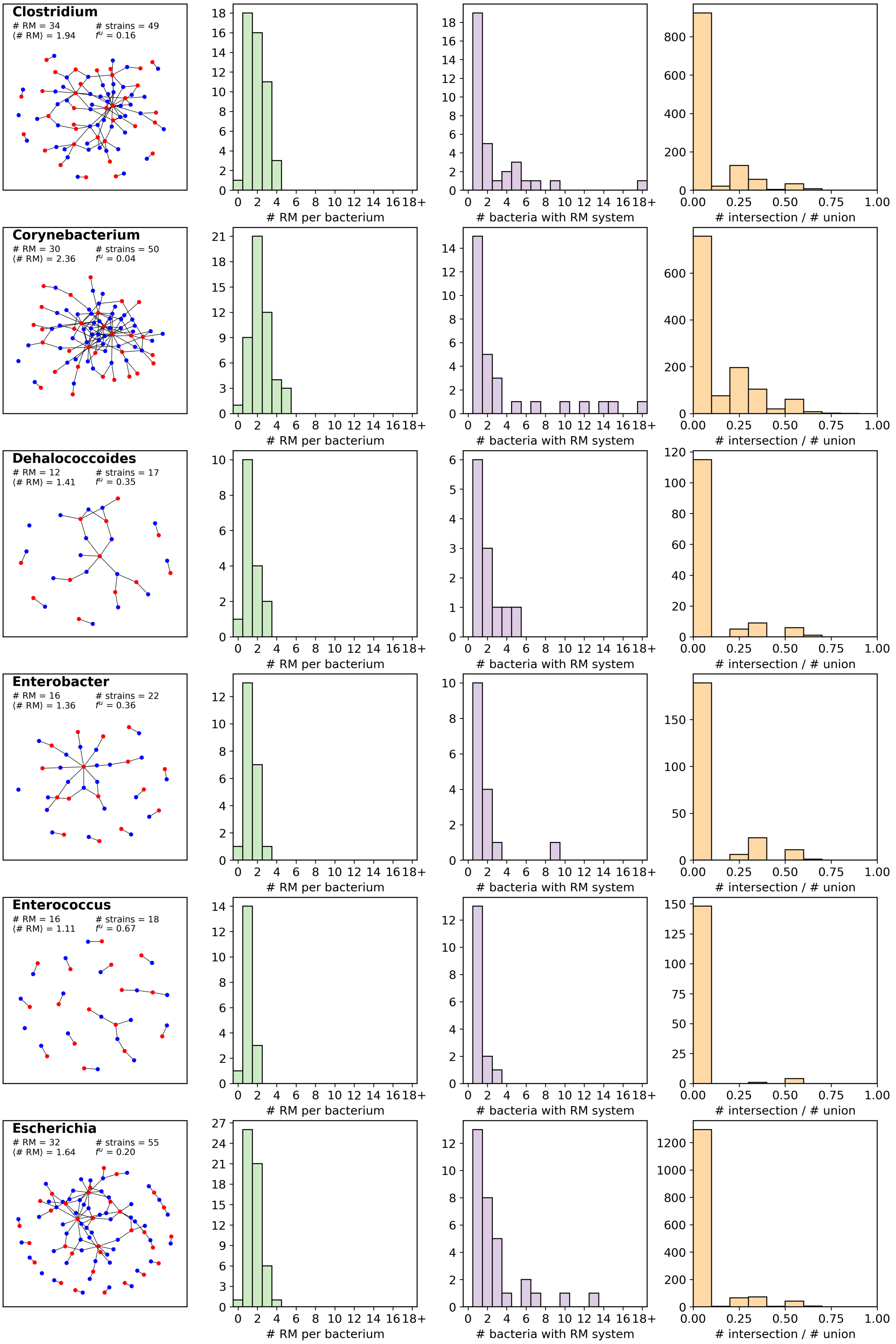


Figure S2 (cont.)

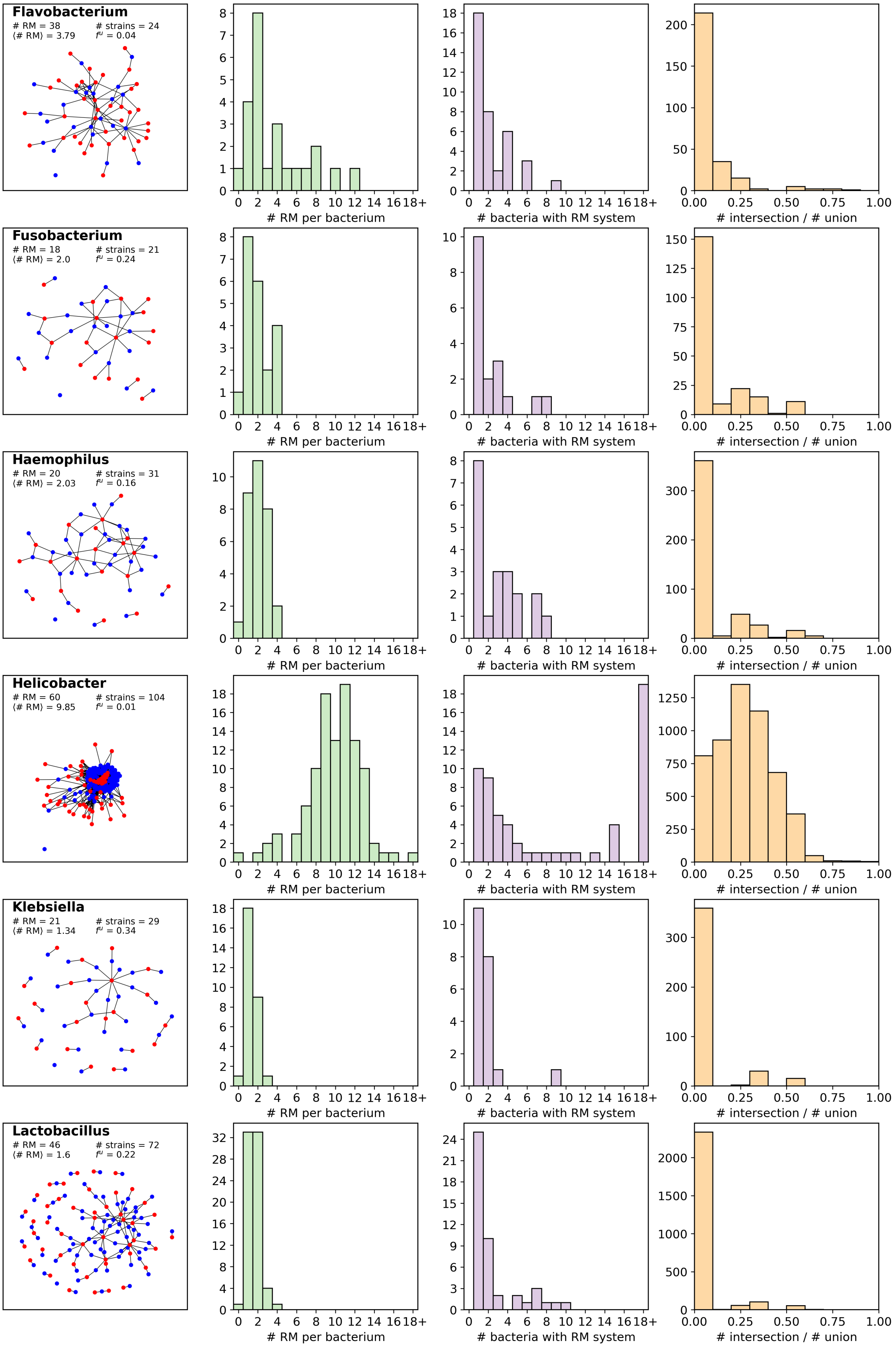


Figure S2 (cont.)

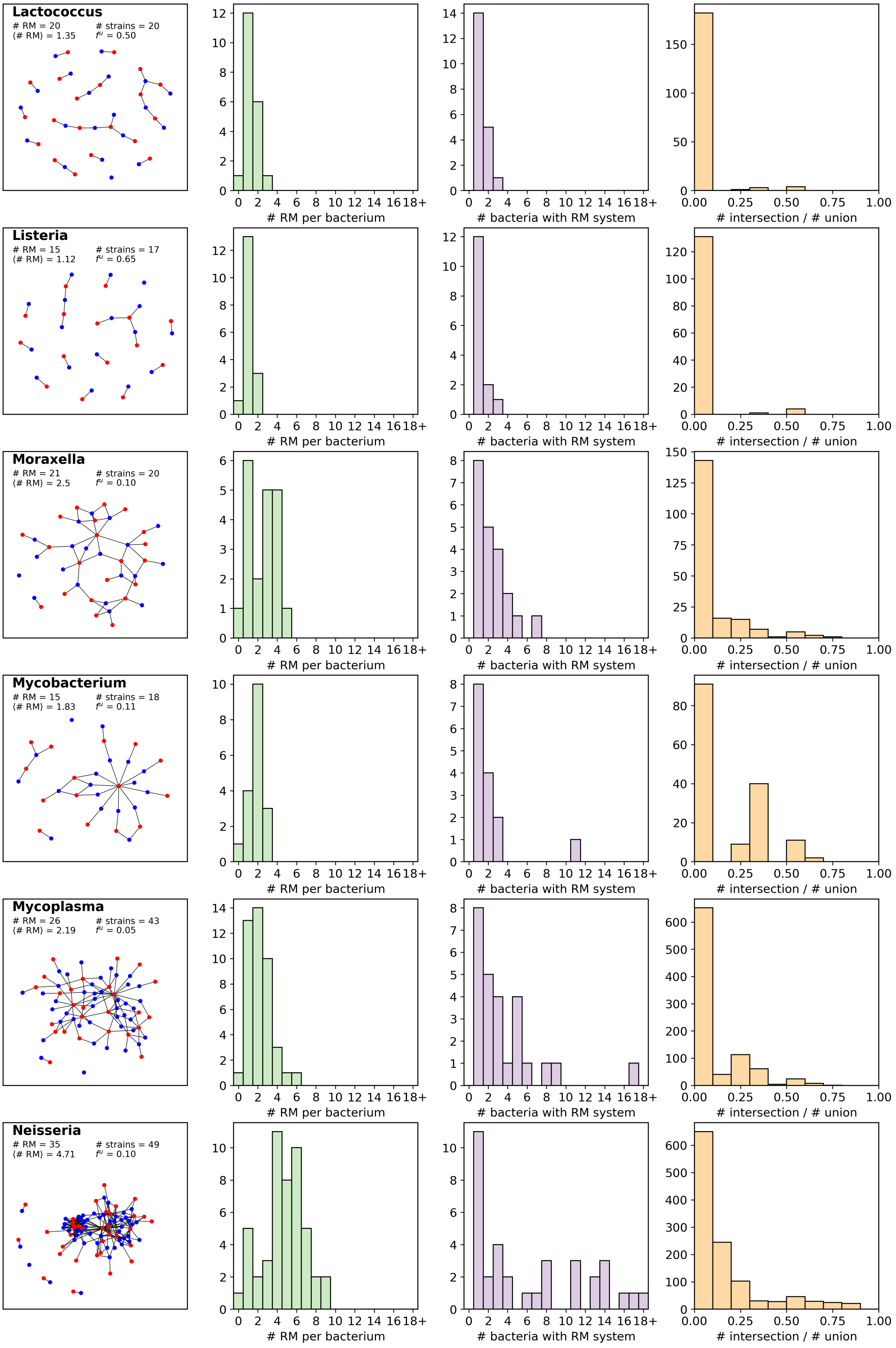


Figure S2 (cont.)

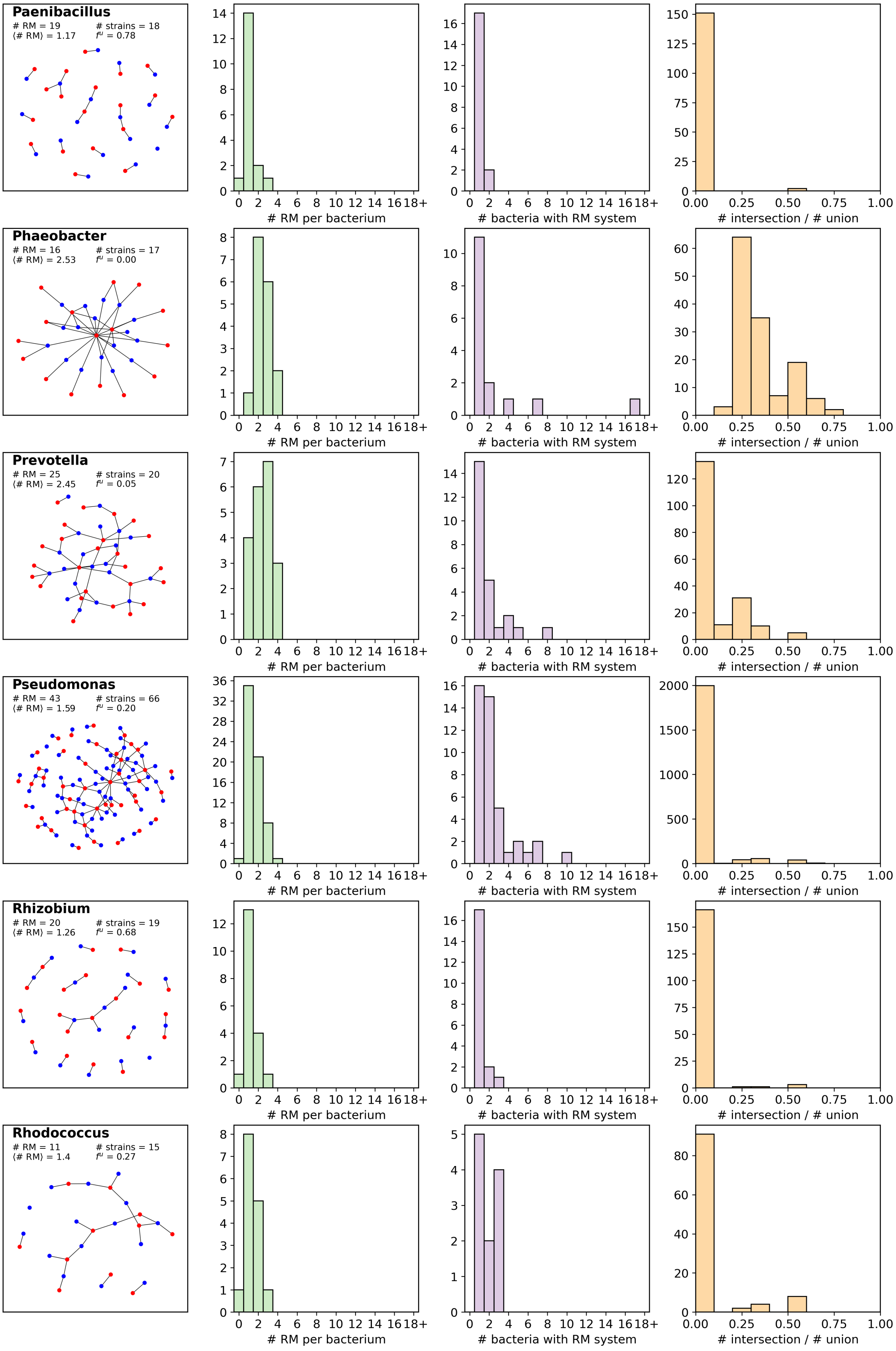


Figure S2 (cont.)

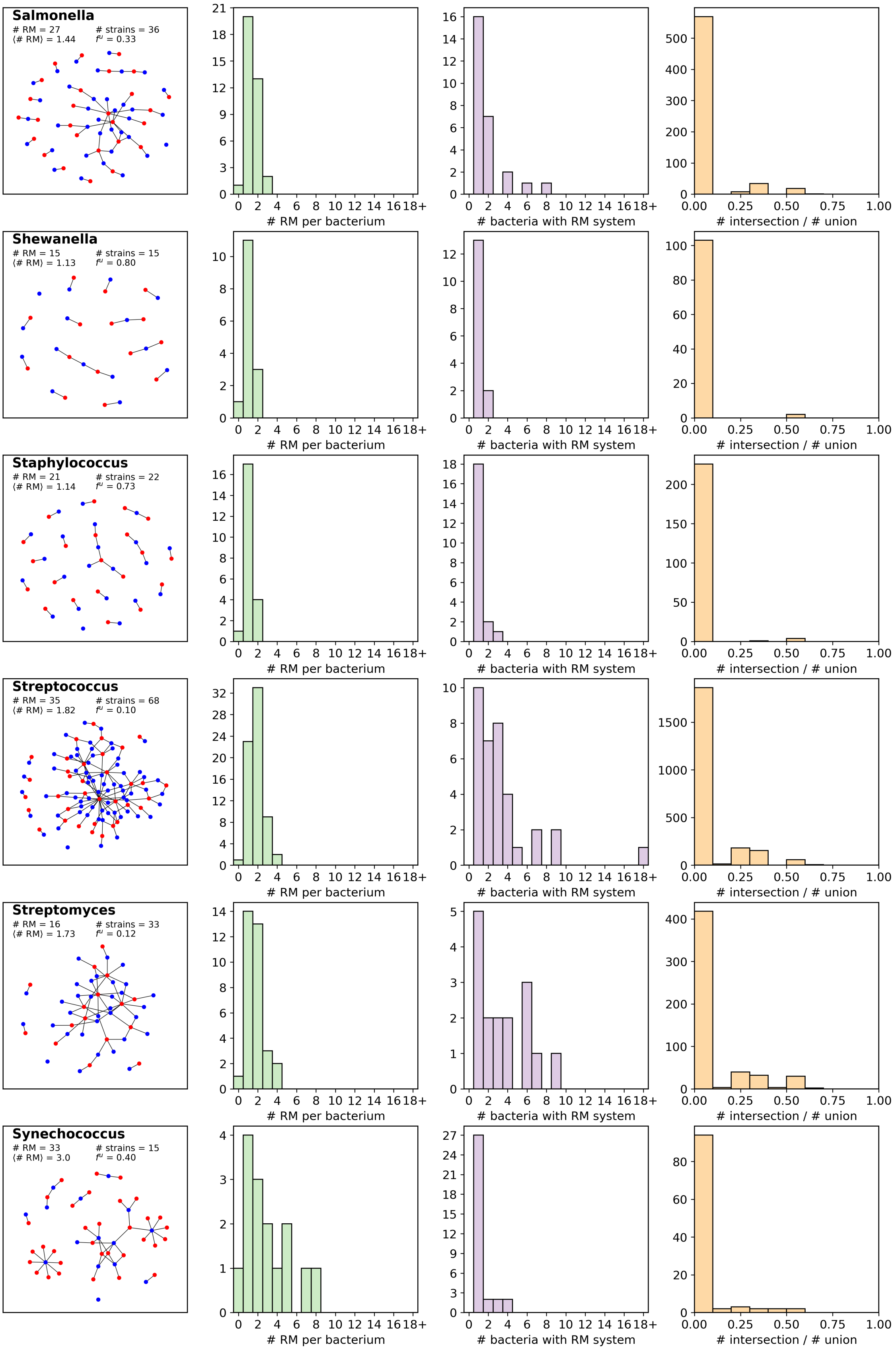


Figure S2 (cont.)

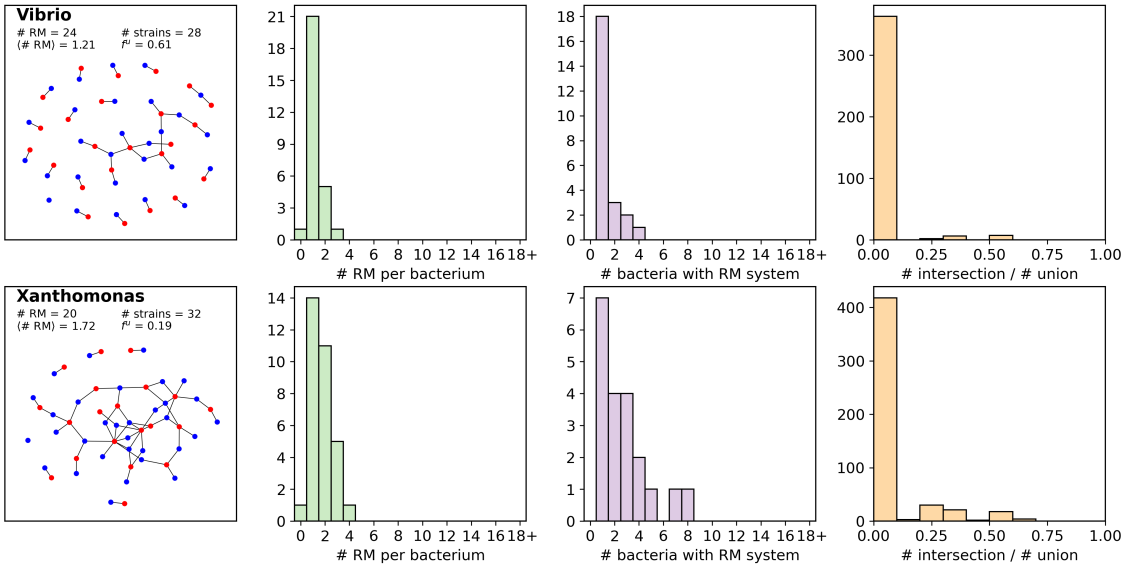


Figure S2 (cont.)

### S4 Comparison with previous experiments

Using a recent analysis of RM genes among *Halobacteria*[3], and data from RM systems in *Salmonella*[2], we compare the distributions of RM systems with our data from NCBI RefSeq[4] and REBASE[1]. In their respective papers, the authors map the presence and absence of RM genes for 217 *Halobacteria* and 221 *Salmonella enterica* subsp. *enterica*. For the *Halobacteria*, the authors identify 26 genes which are related to RM systems as well as 22 "weaker candidates". The salmonella data is more diverse and contains 113 identified RM systems.

For these data sets, we also filter out strains that have the same RM composition. After filtering, we end up with 117 unique *Salmonella* and 200 *Halobacteria*. Which means that we have filtered out 104 and 17 strains respectively.

In figure S3, we plot the distributions of RM systems and the overlap measures for these two data sets (compare with figure 2).

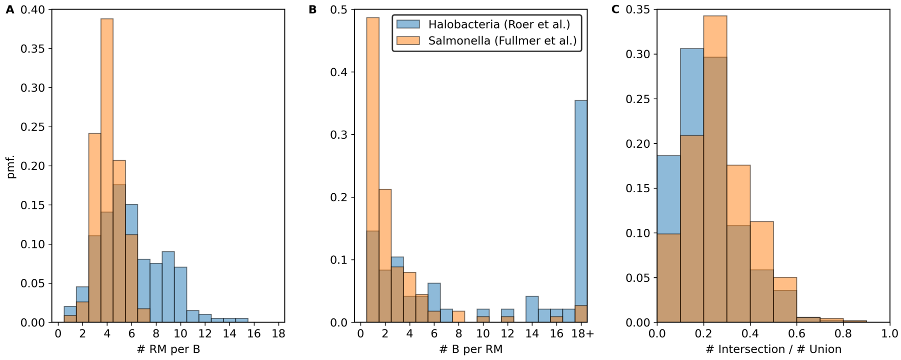


Figure S3: Analysis of RM system distribution in *Halobacteria* (blue bars) and *Salmonella* (orange bars) from the data of [2, 3]. (A) The number of RM systems in the bacteria. (B) The number of bacteria containing a given RM system. (C) The overlap between any combination of two bacteria, defined as the ratio of the number of shared RM system to the number of unique RM systems.

Both distributions show a peak around $\sim 4-5$ RM systems per bacterium but the distribution for the *Halobacteria* is substantially wider than the distribution for Salmonella.

The overlap distribution in both data sets shows a much higher overlap than what we observe for our full data set (compare figures 2B and S3C). The results are similar for both distributions with *Salmonella* leaning towards higher overlap than the *Halobacteria*. For both data sets, roughly

1 in 3 RM systems are found in another random bacteria (if $i$ and $j$ each has three RM systems, then with one overlapping RM system $|S_i \cap S_j| = 1$ while the union will contain $|S_i \cup S_j| = 5$ RM systems).

Next, we compute the RM networks for these data sets (see figure S4). For *Halobacteria*, we find that the RM systems are shared relatively uniformly, while for *Salmonella*, a few RM systems are shared among many strains while many RM systems are shared among only a few strains.

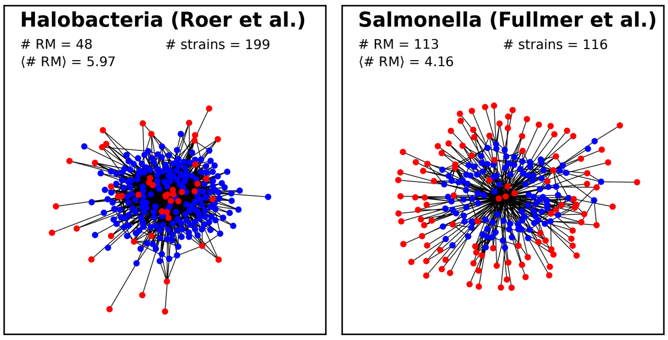


Figure S4: The networks of RM systems, computed as in the main text, for the two data sets: *Halobacteria*[3] and *Salmonella*[2]

#### S5 Breakdown of the overlap distribution

The distributions of the overlap measure, $I$ / $U$, in Fig. S2 also vary between the genera. In Fig. 2E, these distributions have been combined to show the overall overlap distribution, but this figure does not account for the differences in the number of RM systems per strain. In this section, we summarize the overlap measure across the different genera, where we account for these differences in the number of strains. In Fig. S5A we show the overlap between strain $i$ and strain $j$ of the same genus, but we stratify the results by the number of RM systems in the pair of strains.

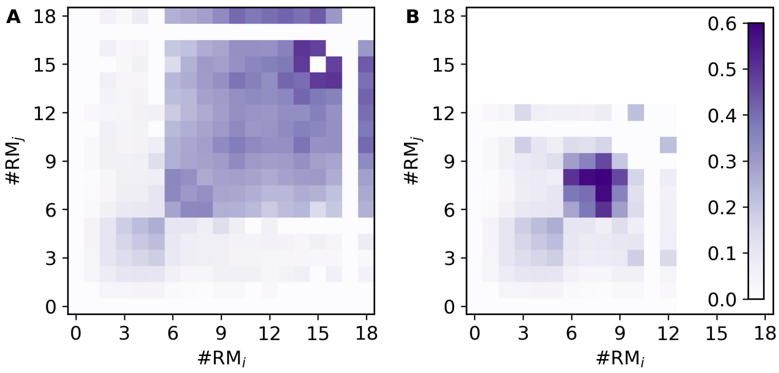


Figure S5: Breakdown of the overlap distribution. (A) the distribution of overlap between all strain-pairs, strain $i$ and strain $j$, within each of the 42 genera stratified by the number of RM systems in each of the strains. (B) similar to (A) but excluding *Helicobacter*.

In doing so, we observe that pairs with high overlap are spread widely in this representation, and are not restricted to strains having either only low number of RM systems. In Fig. S5B, we repeat the analysis without *Helicobacter*. This comparison shows that the structure in the overlap measure for pairs with 6 or more RM systems is dominated by the contribution of *Helicobacter*, which has the highest average number of RM systems per strain.

### Part II
### Model

#### S6 Fitness advantage of restriction site avoidance

Our model implementation allows us to include more than a single phage. These phages use the same parameters, but vary in which RM systems they are immune to. In the model, this translates to each phage modifying the set of effective RM systems:

$$S^k \equiv \{r \in S_i \mid r \notin (I^k \cup S_j)\} \tag{S1}$$

where the index $k$ denotes the phage strain. $r, S_i, S_j$ are as in the main text.

Notice that this does not change the growth rate of the bacteria, since these depend only on $S_i$, but it does change the efficacies of the RM systems:

$$\Omega^k_{i,j} = \prod_{r \in S^k} \omega_r \tag{S2}$$

Mathematically, the model is then described by the equations:

$$\dot{b}_i = \Gamma_i b_i(1 - B/C) - b_i \sum_k \sum_j \eta_j \Omega^k_{i,j} p^k_j - \alpha b_i \tag{S3}$$

$$\dot{p}^k_i = \beta_i b_i \sum_j \eta_j \Omega^k_{i,j} p^k_j - \eta_i p^k_i B - \delta_i p^k_i \tag{S4}$$

With this framework, we simulate a competition assay between two phage strains and three bacterial strains. The bacteria have $K = 2$, and all combinations are included. One of the phage strains is sensitive to all RM systems while the other is immune to one of the RM systems.

We show the results of the simulated assay in Fig. S6. Noticeably, the phage strain that is immune to one RM system does better than its counterpart in all cases and often dominates the other substantially.

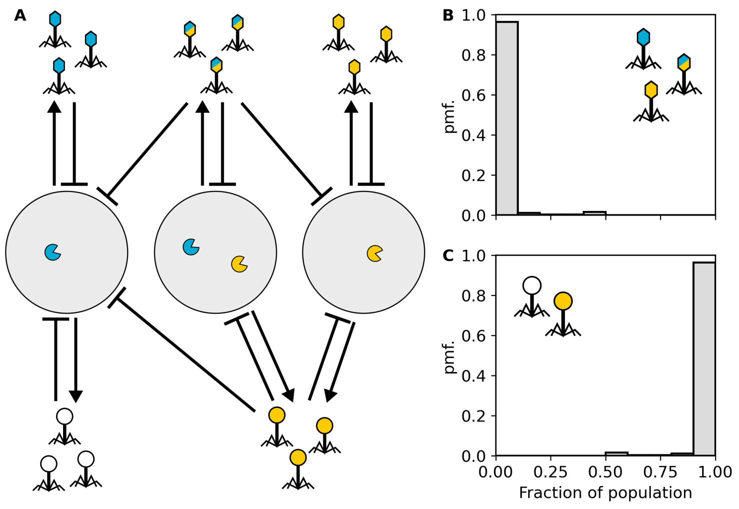


Figure S6: The competitive advantage of restriction site avoidance. We run 250 simulations where two phage strains are mixed with three bacterial strains. (A) Illustration of the model system. The three bacterial strains share two RM systems among them which are labelled by color. One phage variant (hexagon head) contains the recognition sites of both RM systems. The other phage (round head) has lost the recognition site of the blue RM system. (B) and (C) show the relative fraction of each phage variant at the end of the simulations. Simulations deviate from default parameters with $\beta = 25$.

### S7 Possible solutions to the model equations

#### S7.1 Low diversity limit

In this section, we consider in detail the situation from Fig. 5B where there are two available RM systems, A and B, for the bacteria choose from. This means there are three bacterial populations: $b_A$, $b_B$, $b_{AB}$, and the corresponding three epigenetic phage variants: $p_A$, $p_B$, $p_{AB}$. We now assume that each RM system has equal cost and effectiveness ($\gamma = \gamma_A = \gamma_B$ and $\omega = \omega_A = \omega_B$), which gives the follow set of six equations:

$$\dot{b}_{A/B} = \gamma b_{A/B}(1 - B/C) - \eta b_{A/B}\left(p_{A/B} + p_{AB} + \omega p_{B/A}\right) - \alpha b_{A/B} \quad \text{(S5)}$$

$$\dot{b}_{AB} = \gamma^2 b_{AB}(1 - B/C) - \eta b_{AB}\left(p_{AB} + \omega(p_A + p_B)\right) - \alpha b_{AB} \quad \text{(S6)}$$

$$\dot{p}_{A/B} = \beta\eta b_{A/B}\left(p_{A/B} + p_{AB} + \omega p_{B/A}\right) - \eta p_{A/B}B - \delta p_{A/B} \quad \text{(S7)}$$

$$\dot{p}_{AB} = \beta\eta b_{AB}\left(p_{AB} + \omega(p_A + p_B)\right) - \eta p_{AB}B - \delta p_{AB}. \quad \text{(S8)}$$

We solve equations (S5)-(S8) to determine the boundary of the coexistence region. First we consider the steady state $\dot{p}_{A/B} = 0$ in the limiting case where $b_{AB} = p_{AB} = 0$ but $p_{A/B} > 0$:

$$\beta\eta b_{A/B}\left(1 + \omega\right) - \eta B - \delta = 0 \quad \text{(S9)}$$

Since $b_{AB} = 0$, the total bacterial biomass is $B = 2b_{A/B}$, and we can solve $b_{A/B}$ right at the coexistence boundary:

$$b_{A/B} = \frac{\delta}{\eta\left(\beta\left(1 + \omega\right) - 2\right)} \quad \text{(S10)}$$

We next combine the equations $\gamma\dot{b}_{A/B} = 0$ and $\dot{b}_{AB} = 0$ to get:

$$\gamma\eta\left(p_{A/B} + p_{AB} + \omega p_{B/A}\right) + \gamma\alpha = \eta\left(p_{AB} + 2\omega p_{A/B}\right) + \alpha \quad \text{(S11)}$$

If we take the limiting case where $p_{AB}$ approaches 0 and condense the above equation we get:

$$p_{A/B}\left(\gamma\eta\left(1 + \omega\right) - 2\eta\omega\right) = \alpha\left(1 - \gamma\right) \quad \text{(S12)}$$

We next fully isolate $p_{A/B}$:

$$p_{A/B} = \frac{\alpha\left(1 - \gamma\right)}{\eta\left(\gamma\left(1 + \omega\right) - 2\omega\right)} \quad \text{(S13)}$$

Next, we insert equations (S10) and (S13) into $\dot{b}_{A/B} = 0$ in the limit where $b_{AB} = p_{AB} = 0$:

$$\gamma\left(1 - \frac{2\delta}{C\eta(\beta(1 + \omega) - 2)}\right) - \eta\left(1 + \omega\right)\frac{\alpha\left(1 - \gamma\right)}{\eta\left(\gamma\left(1 + \omega\right) - 2\omega\right)} - \alpha = 0 \quad \text{(S14)}$$

Finally, we simplify this expression to determine the boundary of the coexistence region described by the function:

$$g(\gamma, \omega) = \gamma\left(\gamma(1 + \omega) - 2\omega\right)\underbrace{\left(1 - \frac{2\delta}{C\eta(\beta(1 + \omega) - 2)}\right)}_{k} - \alpha(1 - \omega) = 0. \quad \text{(S15)}$$

We now define $\gamma = f(\omega)$, and look for solutions to $g(f(\omega), \omega) = 0$, which leads to the quadratic equation for $f(\omega)$:

$$(1 + \omega)kf(\omega)^2 - 2\omega kf(\omega) - \alpha(1 - \omega) = 0. \quad \text{(S16)}$$

with solutions:

$$f(\omega) = \frac{\omega k \pm \sqrt{\omega^2k^2 + \alpha(1 + \omega)(1 - \omega)k}}{(1 + \omega)k} \quad \text{(S17)}$$

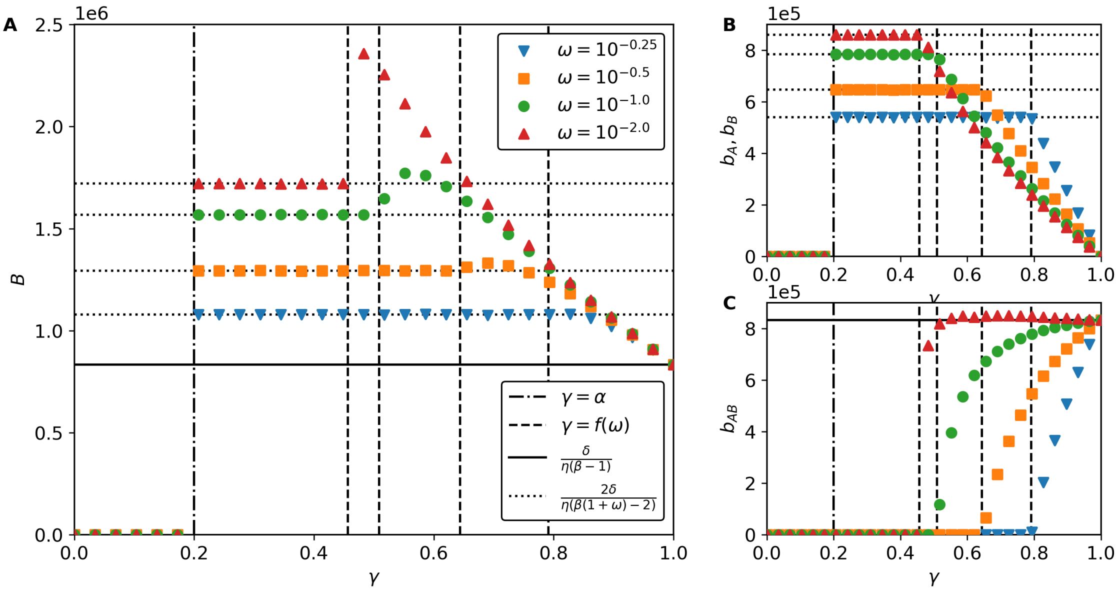


Figure S7: Example solutions with two RM system. (A) Total biomass $B = b_{AB} + b_A + b_B$. (B) Populations of hosts with single RM system. (C) Population of host with both RM systems. Simulations deviate from default parameters with $\beta = 25$ and $T = 10^4$.

In the limit where $\omega = 0$, we have:

$$f(0) = \pm\sqrt{\frac{\alpha}{k}} \tag{S18}$$

If we consider the parameters typically used in phage-bacterial modelling, the factor $k$ is roughly 1 since $\delta$ will be small ($\sim 10^{-2}$), $C\eta$ will be within a few orders of magnitude from unity, and $\beta$ will be relatively large ($\sim 10^2$):

$$k = 1 - \frac{2\delta}{C\eta(\beta - 2)} \approx 1 \tag{S19}$$

In the limit where $\omega = 1$, we have:

$$f(1) = \frac{k \pm k}{2k} \tag{S20}$$

If $\omega$ is zero, i.e., the RM systems are fully perfect, the strain carrying both RM systems can only go extinct if its reduced growth rate $\gamma^2$ is less than $\alpha$, and coexistence requires $f(0) > \sqrt{\alpha}$. When the RM systems are fully inept, that is when $\omega = 1$, the strain carrying both has no benefit from its RM systems and the growth rates must be degenerate for the strains to coexist, meaning that coexistence requires $f(1) = 1$. Combined, this means that the positive solution fits our requirement:

$$f(\omega) = \frac{\omega k + \sqrt{\omega^2 k^2 + k\alpha(1+\omega)(1-\omega)}}{(1+\omega)k}. \tag{S21}$$

In Fig. S7 we show solutions to bacterial populations are possible as we vary the growth rate $\gamma$ and the RM imperfection $\omega$.

Here we can identify regions with possible solutions: When $\gamma < \alpha$ no bacteria can survive as they grow slower than they are being diluted. Additionally, we see that $\gamma$ has to be above some value in order for the $b_{AB}$ population to exist.

When $\alpha < \gamma < f(\omega)$ only the bacteria with a single RM system are able to survive and they settle at a level determined by the effectiveness of the RM system: $b_A = b_B = \delta/(\eta(\beta(1+\omega) - 2))$. In this range of $\gamma$, the host with both RM systems cannot survive since its growth rate, $\gamma^2$, is too small to counteract the dilution and phage predation. For $\gamma$ values above $f(\omega)$ we begin to see coexistence of all species. In most cases, the host with two RM systems will out-compete the other bacteria, leading to a net loss of biomass. This effect gets stronger as $\gamma$ approaches 1, where the difference between $\gamma$ and $\gamma^2$ gets relatively smaller, meaning that the hosts with single RM systems lose their growth advantage. If, however, the RM systems have high effectiveness, $\omega \ll 1$, the total biomass can be substantially higher near the coexistence boundary.

#### S7.2 High diversity limit

In this section we consider a scenario where the bacterial diversity is $D$. In this scenario, we model a background population of bacterial strains which have unique RM systems, and a number of triplets $T$ which are each linked by their RM systems. In particular, we discuss two types of triplets which we denote "hierarchical tripelets" and "looped triplets". "hierarchical triplets" are of the type we just discussed: triplets of the $A/B/AB$ motif, whereas the "looped triplets" are of the $AB/BC/AC$ motif. We solve in detail the "hierarchical triplets" and summarise the solution to the "looped triplets" system which follows the same steps.

##### S7.2.1 Solution to embedded hierarchical triplets

In this section, we consider in detail the situation where there are $T$ independent hierarchical triplets, $(A, B, AB)$-like, in an ecosystem with a total of $D$ strains. Each triplet is independent, meaning the RM systems it contains are not present elsewhere in the ecosystem. The "background" strains $b_i$ are all assumed to have a unique RM system. This means there are bacterial populations: $b_i$, $b_1$, and $b_2$. Here $b_1$ is the biomass of two strains in the triplets with a single RM system, and $b_2$ is the biomass of the strain in the triplet with two RM ssytems. The corresponding epigenetic phage variants have populations: $p_i$, $p_1$, and $p_2$. We now assume that each RM system has equal cost and effectiveness, which gives the follow set of 2D equations:

$$\dot{b}_i = \gamma b_i(1 - B/C) - \eta b_i \left(p_i + \omega \left[(D - 3T - 1)p_i + 2Tp_1 + Tp_2\right]\right) - \alpha b_i \tag{S22}$$

$$\dot{b}_1 = \gamma b_1(1 - B/C) - \eta b_1 \left(p_1 + p_2 + \omega \left[(D - 3T)p_i + (2T - 1)p_1 + (T - 1)p_2\right]\right) - \alpha b_1 \tag{S23}$$

$$\dot{b}_2 = \gamma^2 b_2(1 - B/C) - \eta b_2 \left(p_2 + 2\omega p_1 + \omega^2 \left[(D - 3T)p_i + (2T - 2)p_1 + (T - 1)p_2\right]\right) - \alpha b_2 \tag{S24}$$

$$\dot{p}_i = \beta\eta b_i \left(p_i + \omega \left[(D - 3T - 1)p_i + 2Tp_1 + Tp_2\right]\right) - \eta p_i B - \delta p_i \tag{S25}$$

$$\dot{p}_1 = \beta\eta b_1 \left(p_1 + p_2 + \omega \left[(D - 3T)p_i + (2T - 1)p_1 + (T - 1)p_2\right]\right) - \eta p_1 B - \delta p_1 \tag{S26}$$

$$\dot{p}_2 = \beta\eta b_2 \left(p_2 + 2\omega p_1 + \omega^2 \left[(D - 3T)p_i + (2T - 2)p_1 + (T - 1)p_2\right]\right) - \eta p_2 B - \delta p_2. \tag{S27}$$

As before, we solve equations (S22)-(S27) to determine the boundary of the coexistence region. First we consider the steady state $\dot{p}_1 = 0$ in the limiting case where $b_2 = p_2 = 0$ but $p_1 > 0$. Since the strains with two RM systems have zero biomass, the single strains are all equal ($b = b_i = b_1$, $p = p_i = p_1$):

$$\beta\eta b \left(p + \omega \left[(D - 3T - 1)p + 2Tp\right]\right) = \eta p B + \delta p \tag{S28}$$

Since $b_2 = 0$, the total bacterial biomass is $B = (D-T)b$, and we can solve $b$ right at the coexistence boundary:

$$\beta\eta b \left(1 + \omega \left[(D - 3T - 1) + 2T\right]\right) = \eta(D - T)b + \delta \tag{S29}$$

$$b = \frac{\delta}{\eta \left(\beta \left(1 + \omega \left[(D - T - 1)\right]\right) - (D - T)\right)} \tag{S30}$$

We next combine the equations $\gamma\dot{b} = 0$, $\gamma\dot{b_1} = 0$ and $\dot{b}_2 = 0$ to get:

$$\gamma\eta \left(p + \omega \left[(D - 3T - 1)p + 2Tp_1 + Tp_2\right]\right) - + \gamma\alpha \tag{S31}$$

$$= \gamma\eta \left(p_1 + p_2 + \omega \left[(D - 3T)p + (2T - 1)p_1 + (T - 1)p_2\right]\right) + \gamma\alpha \tag{S32}$$

$$= \eta \left(p_2 + 2\omega p_1 + \omega^2 \left[(D - 3T)p + (2T - 2)p_1 + (T - 1)p_2\right]\right) + \alpha \tag{S33}$$

If we take the limiting case where $p_2$ approaches 0 and condense the above equation we get:

$$\gamma\eta\left(p+\omega\left[(D-3T-1)p+2Tp\right]\right)+\gamma\alpha \tag{S34}$$
$$=\gamma\eta\left(p+\omega\left[(D-3T)p+(2T-1)p\right]\right)+\gamma\alpha \tag{S35}$$
$$=\eta\left(2\omega p+\omega^2\left[(D-3T)p+(2T-2)p\right]\right)+\alpha \tag{S36}$$

$$\eta p\left[\gamma\left(1+\omega(D-T-1)\right)-\left(2\omega+\omega^2(D-T-2)\right)\right]=\alpha(1-\gamma) \tag{S37}$$

We next fully isolate $p$:

$$p=\frac{\alpha(1-\gamma)}{\eta\left[\gamma\left(1+\omega(D-T-1)\right)-\left(2\omega+\omega^2(D-T-2)\right)\right]} \tag{S38}$$

Next, we insert equations (S30) and (S38) into $\dot{b}_1=0$ in the limit where $b_2=p_2=0$. But first we define the following quantities:

$$k_1=1+\omega(D-T-1) \tag{S39}$$
$$k_2=2\omega+\omega^2(D-T-2) \tag{S40}$$

$$\gamma\left(1-\frac{(D-T)\delta/C}{\eta\left[\beta k_1-(D-T)\right]}\right)-\alpha-\eta k_1 p=0 \tag{S41}$$
$$\gamma\left(1-\frac{(D-T)\delta/C}{\eta\left[\beta k_1-(D-T)\right]}\right)-\alpha-\eta k_1\frac{\alpha(1-\gamma)}{\eta\left[\gamma k_1-k_2\right]}=0 \tag{S42}$$
$$\tag{S43}$$

Finally, we simplify this expression to determine the boundary of the coexistence region described by the function:

$$g(\gamma,\omega)=\gamma\left(\gamma k_1-k_2\right)\underbrace{\left(1-\frac{(D-T)\delta/C}{\eta\left[\beta k_1-(D-T)\right]}\right)}_{k_3}-\alpha\left(\gamma k_1-k_2\right)-k_1\alpha(1-\gamma)=0 \tag{S44}$$

We now define $\gamma=f(\omega)$, and look for solutions to $g(f(\omega),\omega)=0$, which leads to the quadratic equation for $f(\omega)$:

$$k_1k_3f(\omega)^2-k_2k_3f(\omega)-\alpha(k_2-k_1)=0. \tag{S45}$$

with solutions:

$$f(\omega)=\frac{k_1k_2\pm\sqrt{k_1^2k_2^2-4\alpha k_1k_3(k_2-k_1)}}{2k_1k_3} \tag{S46}$$

From here, the solution follows the same arguments as above and the positive solution can be identified.

##### S7.2.2 Solution to embedded looped triplets

In this section, we consider in detail the situation where there are $T$ independent triplets $(AB, BC, AC)$-like in an ecosystem with a total of $D$ strains. Each triplet is independent, meaning the RM systems it contains are not present elsewhere in the ecosystem. The "background" strains $b_i$ are all assumed to have a unique RM system. This means there are bacterial populations: $b_i$ and $b_2$. Here $b_2$ are the strains in the triplets with two RM system (i.e., all three strains in the triplet). The systems have the corresponding epigenetic phage variants whose populations are denoted $p_i$ and $p_2$. We now assume that each RM system has equal cost and effectiveness, which gives the follow set of 2D equations:

$$\dot{b}_i=\gamma b_i(1-B/C)-\eta b_i\left(p_i+\omega\left[(D-3T-1)p_i+3Tp_2\right]\right)-\alpha b_i \tag{S47}$$
$$\dot{b}_2=\gamma^2 b_2(1-B/C)-\eta b_2\left(p_2+2\omega p_2+\omega^2(D-3T)p_i\right)-\alpha b_2 \tag{S48}$$
$$\dot{p}_i=\beta\eta b_i\left(p_i+\omega\left[(D-3T-1)p_i+3Tp_2\right]\right)-\eta p_i B-\delta p_i \tag{S49}$$
$$\dot{p}_2=\beta\eta b_2\left(p_2+2\omega p_2+\omega^2(D-3T)p_i\right)-\eta p_2 B-\delta p_2. \tag{S50}$$

Solving the equations leads to the factors

$$k_1 = 1 + \omega(D - 3T - 1) \tag{S51}$$

$$k_2 = \omega^2(D - 3T) \tag{S52}$$

$$k_3 = -\frac{(D - 3T)\delta/C}{\eta\,[\beta k_1 - (D - 3T)]} \tag{S53}$$

##### S7.2.3 Embedded triplets

With the solution for $f(\omega)$ for the above scenarios, we can plot the parameter regions where the triplets are able to coexist with the single RM strain bacteria. In Fig. S8 we show these demarcation lines as we vary the number of triplets $T$ in a population of $D = 100$ bacterial strains.

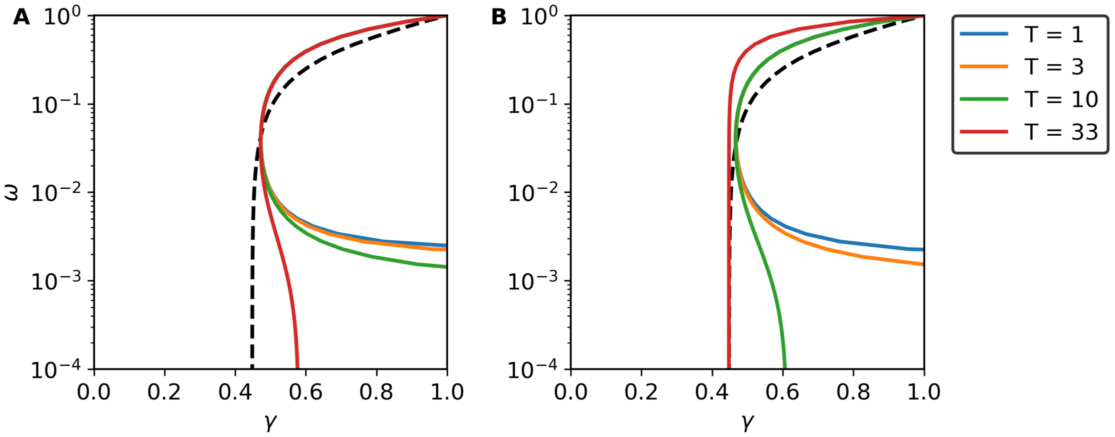


Figure S8: Triplet coexistence in the high diversity limit. We show the solutions to $f(\omega)$ when $D = 100$ for different number of triplets $T$ of (A) hierarchical type and (B) looped type. Black dashed lines in (A) and (B) indicate the solution for a single hierarchical triplet S7.1 in the absence of background unique strains.

We contrast these demarcation lines with the reference scenario of a single hierarchical triplet and no other strains (section S7.1). The solution highlights an interesting restriction on values of $\omega$ that yields viable triplets when the ecosystem predominantly consists of bacterial strains each having a single, but unique, RM system. Here the triplets cannot exist if the value of $\omega$ is sufficiently small. We speculate that this is explained by the lower phage pressure in the high diversity limit (many phages are lost to infecting strains with effective RM systems) which leads to the individual RM systems being strong enough to sufficiently negate the phage pressure. As the number of triplets increase, the demarcation lines shifts, and a wider range of parameters lead to the ecosystem supporting the triplets. Interestingly, for the same value of $D$ and $T$, an ecosystem of looped triplets is supported by a wider range of parameters than an ecosystem of hierarchical triplets, which suggests that increased overlap between bacterial strains is competitively beneficial.

#### S8 Examples of the open-ecosystem

In this section we show examples runs for the simulations of the open-ecosystem where we vary number of available RM systems, $K$, to choose from. In Figs. S9(A-C) we include $K = \infty$, $K = 800$ and $K = 50$ RM systems respectively. Since the bacteria can have any combination of these RM systems, this corresponds to roughly $10^{240}$ combinations when $K = 800$ and $10^{15}$ combinations when $K = 50$, each with their own $(\Gamma_i, \Omega_i)$ values.

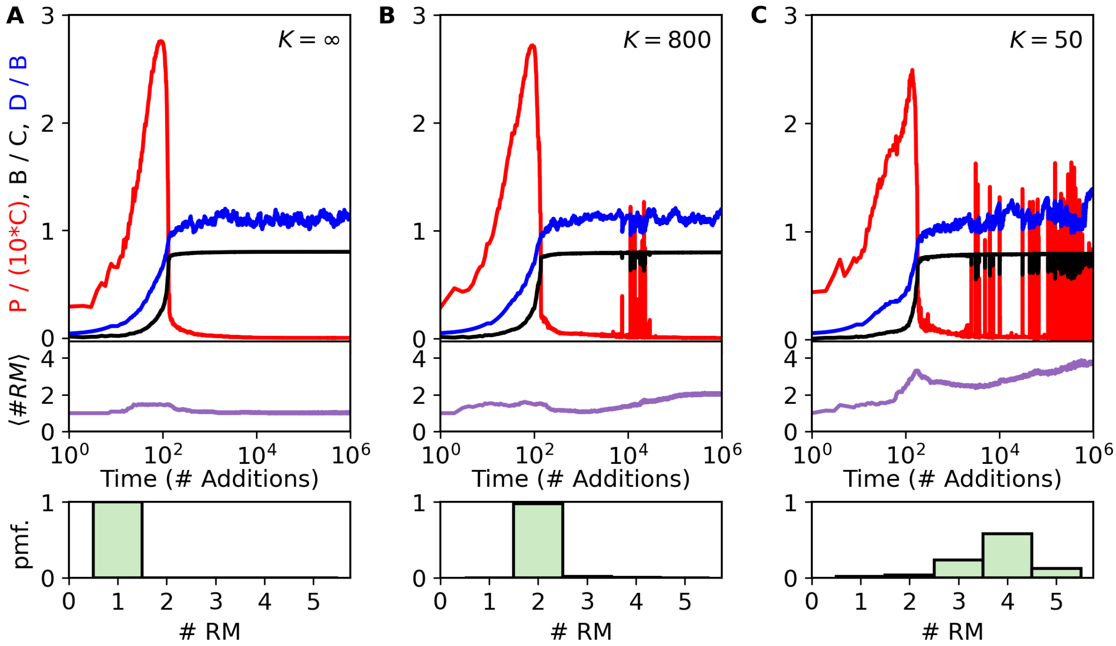


Figure S9: The population dynamics of open ecosystems. Here, every $T = 10^3$ generations, we add a new bacterial strain which carries a novel combination of the $K$ possible RM systems. (A) $K = \infty$, (B) $K = 800$, and (C) $K = 50$. We show the total bacterial density, $B$, in units of the carrying capacity $C$ (black curve), the total phage density $P$ (red curve) in units of $10 \cdot C$, the diversity $D$ (blue curve) in units of the phage burst size $\beta$, and the average number of RM systems per strain $\langle \#RM \rangle$ in the system over time. (D-F) Distributions of the number RM systems each bacterial strain carries at the end of the simulation.

#### S8.1 Triplet composition

Based on section S7.2, we expect the number of triplets to increase over time in our open ecosystem. In Fig. S10(A-B), we show that this behavior is observed, with the number of both hierarchical and looped triplets increasing over time. The number of triplets is larger for smaller repertoires of RM systems (smaller values of $K$) which matches the expected increased overlap in these scenarios. In fig. S10C, we show the fraction of triplets that have the looped motif, and we observe that this fraction is also increasing over time – again matching the expectations laid out in section S7.2.

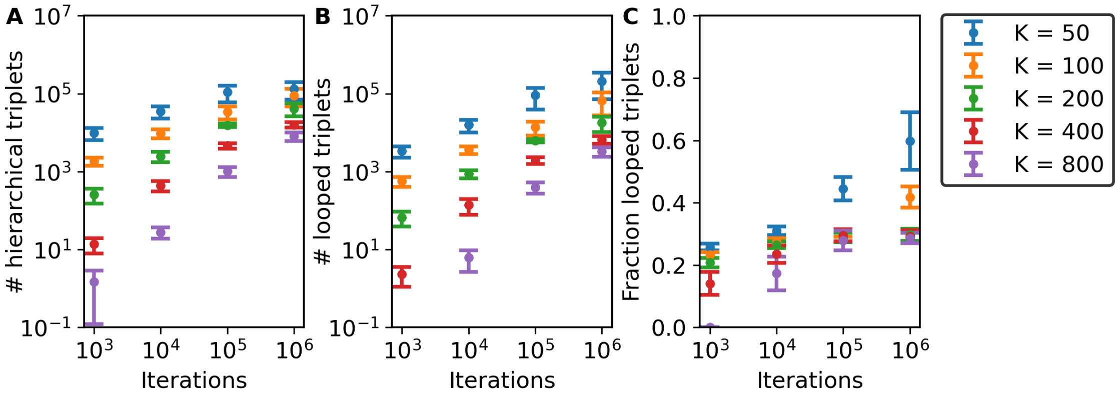


Figure S10: Composition of triplets over time in open ecosystems. We here show: (A) the number of hierarchical triplets, (B) the number of looped triplets, and (C) the fraction of triplets that are looped.

#### S8.2 Phage resurgences

As the number of possible RM systems is restricted (lower values for $K$), the dynamics between the phage and bacteria become increasingly noisy. This is especially apparent in the intermittent periods where the phage population rises from a very small density to (temporarily) be the dominant population in the ecosystem. In Fig. S11 we focus on these noisy dynamics of the open-ecosystem with $K = 50$.

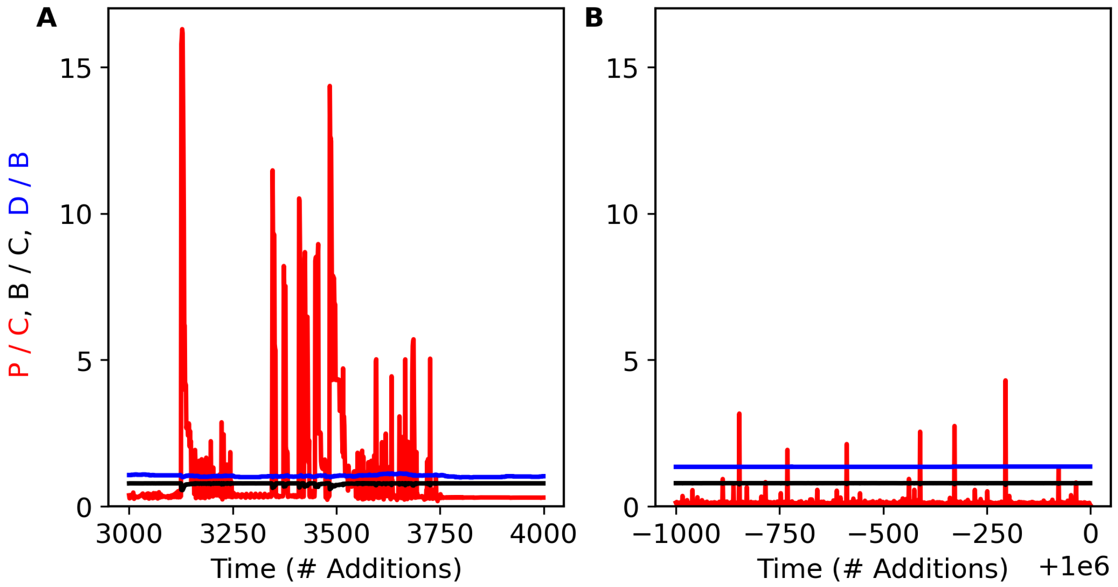


Figure S11: Transient resurgence of phage population in open ecosystems. (A) early phage resurgences: Time-steps 3000 – 4000 (B) later phage resurgences : last 1000 time steps.

From this figure, the structure of the phage resurgences are more clearly visible. In both cases we observe that the phage resurgences are temporary, but that the resurgences early on seem to persist for longer than the later resurgences.

##### S8.3 Extinction curve

In Ref. [5], the authors show that there exists a demarcation line in the $\gamma - \omega$ space which separates strains that will be able to coexist in the ecosystem from those that will not. The authors denote this demarcation line as the "extinction curve" and its location depends on the composition of the ecosystem. In short, successive additions of bacterial strains, in the open ecosystem simulation, move the extinction curve towards higher values of $\gamma$, corresponding to the competition between bacteria selecting for faster growth rates.

The extinction curve is described by the equation:

$$f(\gamma_i, \omega_i) = \gamma_i(1 - B/C) - \alpha - \eta\omega_i P > 0. \tag{S54}$$

Our model is a generalization of their work, and here we apply this concept of the extinction curve to our extended model. In Fig. S12(A-B), we plot the values for $\gamma$ versus the naive value for $\omega$ of all the generated strains after $10^4$ addition of bacterial strains. By "naive" we here specifically don't account for the cross-methylation of the phages in the ecosystem:

$$\omega_i^{naive} = \prod_{r \in S_i} \omega_r.$$

For lower values of $K$, we expect substantial sharing of RM system among the bacteria, and we must therefore represent the efficacy of the RM systems in a manner that accounts for the composition of the ecosystem. In Fig. S12(C-D) we instead use an "effective" $\omega$ for each strain that is computed from the generalized efficacies: $\Omega_{i,k}$ (excluding the self-terms $\Omega_{i,i}$) weighted by the phage population at the end of the simulation. Notice, however, that by weighting by the final phage population, we do not account for the composition when the strains were eliminated.

As expected, we see that the extinction curve is a good predictor for survival when the value of $K$ is large, and we see only small differences between the naive and effective $\omega$s due to the limited overlap of RM systems between bacterial strains. For low values of $K$, the situation is very different. With the naive rates, the extinction curve is a bad predictor for whether bacterial strains will go extinct. However, when we consider the effective $\omega$s, the situation changes and the extinction curve becomes a much better predictor for the survival of the bacterial strains.

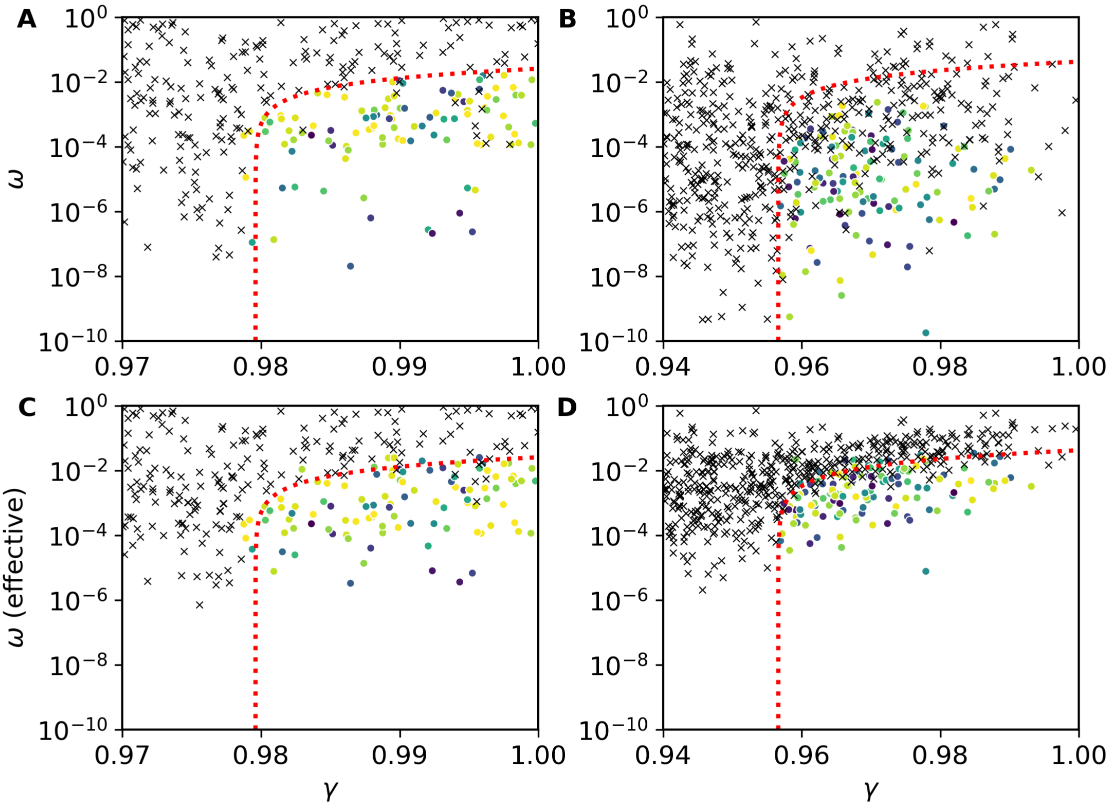


Figure S12: Extinction curve. We show the $\gamma$ and $\omega$ values for all bacterial strains that were added during a simulation with $10^4$ time-steps. The black crosses denote bacterial strains that were eliminated in the simulations and the coloured circles denoting the strains alive at the end of the simulation. The extinction curved based on the final composition of the ecosystem is shown as the dotted red line. (A) $K = 800$. (B) $K = 50$. (C-D) same as (A-B) respectively but with the effective $\omega$ instead of the naive $\omega$ (see text).

#### S9 Parameter values used in simulations

The simulations we have done for this paper use the parameter values listed in table S3 as the default parameters, and any deviation is explicitly specified when applicable.

| Name | Value | Units | Description | References |
|---|---|---|---|---|
| C | $10^8$ | bacteria | Carrying capacity of the ecosystem | |
| $\tau$ | - | hours | Minimal attainable generation time for bacteria | |
| $\eta$ | $10^{-8}$ | $1/\tau$ | Adsorption rate of the phages | [6] assuming $\tau \sim 0.5$ h |
| $\alpha$ | 0.2 | $1/\tau$ | Dilution rate of the ecosystem | |
| $\beta$ | 100 | phages | Burst size of the phages | [6] |
| $\delta$ | 0.2 | $1/\tau$ | Decay rate of the phages | |
| $T$ | $10^3$ | $\tau$ | Interval between bacterial additions | |
| $M$ | 5 | strains | Minimum number of strains in the ecosystem | |

Table S3: Default parameter values used in the simulations.
